# Fascin structural plasticity mediates flexible actin bundle construction

**DOI:** 10.1101/2024.01.03.574123

**Authors:** Rui Gong, Matthew J. Reynolds, Keith R. Carney, Keith Hamilton, Tamara C. Bidone, Gregory M. Alushin

## Abstract

Fascin crosslinks actin filaments (F-actin) into bundles that support tubular membrane protrusions including filopodia and stereocilia. Fascin dysregulation drives aberrant cell migration during metastasis, and fascin inhibitors are under development as cancer therapeutics. Here, we use cryo-electron microscopy, cryo-electron tomography coupled with custom denoising, and computational modeling to probe fascin’s F-actin crosslinking mechanisms across spatial scales. Our fascin crossbridge structure reveals an asymmetric F-actin binding conformation that is allosterically blocked by the inhibitor G2. Reconstructions of seven-filament hexagonal bundle elements, variability analysis, and simulations show how structural plasticity enables fascin to bridge varied inter-filament orientations, accommodating mismatches between F-actin’s helical symmetry and bundle hexagonal packing. Tomography of many-filament bundles and modeling uncovers geometric rules underlying emergent fascin binding patterns, as well as the accumulation of unfavorable crosslinks that limit bundle size. Collectively, this work shows how fascin harnesses fine-tuned nanoscale structural dynamics to build and regulate micron-scale F-actin bundles.

## Introduction

Subcellular cytoskeletal networks are built by filament crosslinking proteins, which specify diverse network geometries to fulfill the cytoskeleton’s myriad functions by poorly defined mechanisms^1–4^. Here, we focus on co-linear actin filament (F-actin) assemblies with uniform polarity (hereafter referred to as “bundles”), which mediate the generation and maintenance of rod-like plasma membrane protrusions that are critical for hearing and balance (stereocilia), nutrient absorption (microvilli), and directed cell migration (filopodia)^5–9^. Filopodia are localized at the leading edge of migrating cells, where they function as dynamic antennae that sense and respond to external cues that instruct cytoskeletal dynamics^8,10–14^. Physiologically, filopodia are necessary for axonal outgrowth and pathfinding, embryonic development, and wound healing^8,11,12,15–20^. Pathologically, they are prominently associated with enhanced migration of metastatic cancer cells^21–25^. While the architecture, composition, and signaling functions of filopodia have been extensively studied, the protein structural mechanisms mediating their assembly and regulation remain poorly understood.

The 55 kDa F-actin crosslinking protein fascin is critical for biogenesis of filopodia, where it is highly abundant, as well as maintenance of stereocilia^26–28^. Crystallographic structures of fascin in the absence of F-actin coupled with saturation mutagenesis studies have led to the proposal that the protein features at least two actin-binding sites (ABS) within the same polypeptide chain, enabling it to function as a monomeric crosslinker^29–32^. Fascin is a clinical biomarker for metastatic cancer, and cells overexpressing fascin feature an overabundance of filopodia-like protrusions that mediate enhanced tissue invasion and migration, which are correlated with high metastatic potential and poor prognosis^33–40^. Protein Kinase C (PKC) phosphorylation on serine 39 negatively regulates fascin, resulting in reduced F-actin binding and bundling and concomitant filopodia disassembly by unknown mechanisms^28,41^. The potent effects of fascin upregulation has motivated the development of small-molecule fascin inhibitors, which have shown promise as cancer therapeutics in preclinical mouse models and are undergoing phase 2 clinical trials for treatment of gynecological and breast cancers^42–44^. Nevertheless, the detailed structural mechanisms enabling fascin to bridge filaments are unknown, limiting insights into its regulation and potential avenues for inhibitor optimization.

Fascin crosslinked actin bundles have been a model system for theoretical and experimental studies of cytoskeletal network assembly principles for decades^31,45–49^. *In vitro*, fascin induces the formation of paracrystalline hexagonal arrays with packing similar to that observed in the cores of sea urchin and mouse filopodia, as well as Drosophila bristles, where fascin is localized i*n vivo*^32,46,50–56^. Early studies of sea urchin fascin suggested that the actin filaments composing these arrays are in perfect register, with identical rotational phase^46,50,52,53^ (i.e., requiring no rotation around their helical axes to superimpose neighboring filaments). Geometric analysis predicted each filament in a bundle would feature three pairs of fascin crossbridges evenly distributed across every axial repeat-length (approximately 13 actin subunits), oriented along the three diagonals of the hexagonal lattice. In this bundle architecture, fascin crossbridges aligned along the same diagonals form linear transverse crossbands perpendicular to the co-linear filament axes (Figures 1A-1C)^46^. However, mismatches between F-actin’s helical symmetry and the translational symmetry of the hexagonal array renders these bridging positions non-identical, leading to speculation that fascin could flex to accommodate geometric strain or modulate F-actin structure^52^. More recent studies of arrays crosslinked by vertebrate fascin *in vitro* and cryo-electron tomography studies of mouse neuronal growth cone filopodia *in situ* have instead suggested noncoherent filament rotational phases, with bundles featuring slanted or chevron-shaped crossbands, implying a more complex pattern of crossbridges (Figure 1B)^31,54,55,57^. Moreover, while arrays with perfect filament registration could theoretically grow indefinitely, bundles within filopodia and those reconstituted from F-actin and vertebrate fascin both feature a maximum of approximately 20-30 laterally-associated filaments^31,49,54,55^, a constraint which maintains filopodial mechanical compliance to support dynamic sampling of the extracellular environment^45^. The mechanisms by which fascin detects and modulates the architecture of F-actin bundles to facilitate filopodia size control have yet to be determined.

**Figure 1.**
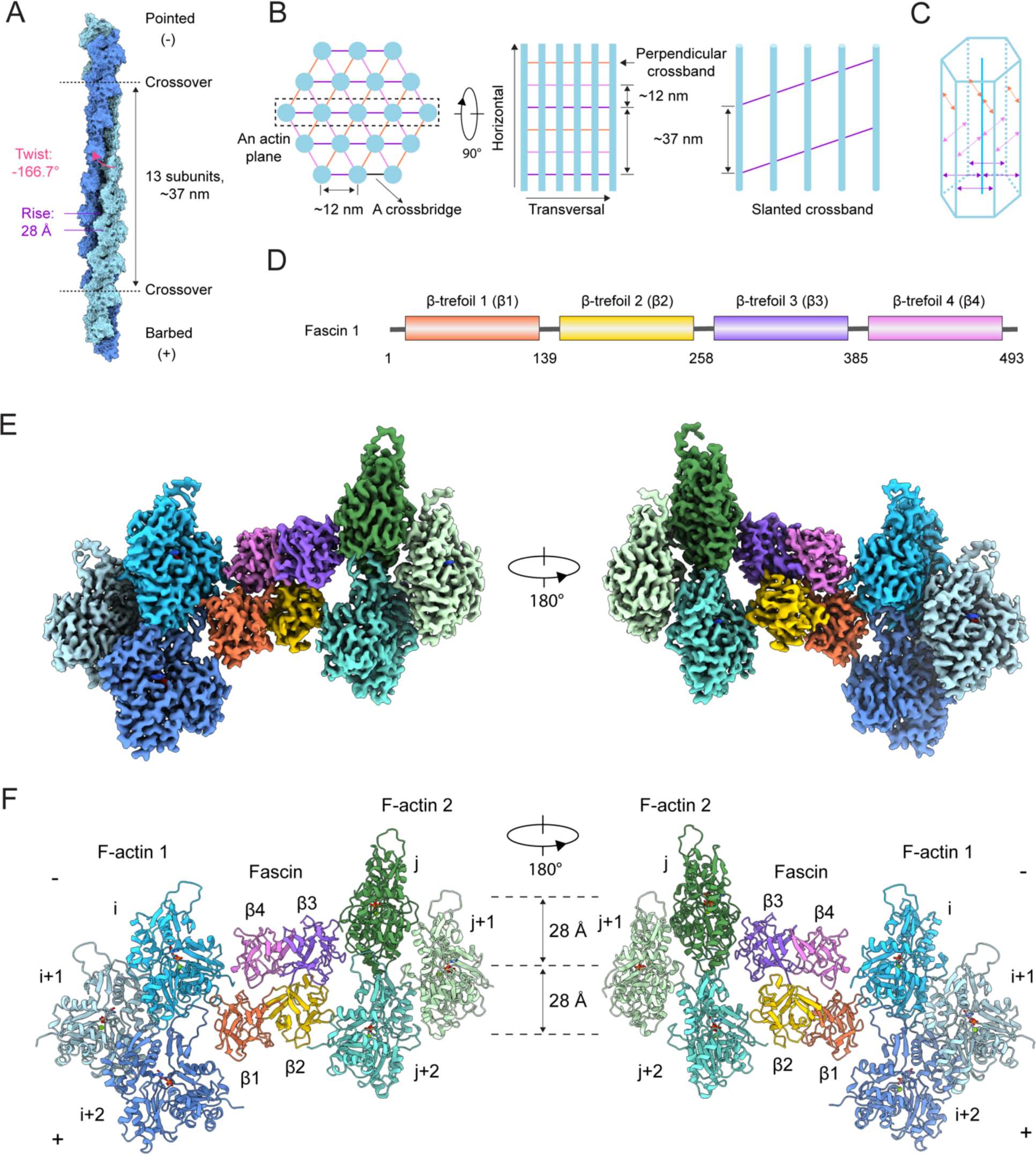
Cryo-EM structure of fascin crosslinking actin filaments. (A) Diagram of F-actin’s helical architecture. Model generated from PDB 76DV. (B) Left: Schematic of end-on and side views of a fascin crosslinked F-actin array. Actin filaments are shown in blue, and fascin crossbridges at each diagonal are depicted as colored lines. Right: Side view of an actin plane (box in left panel) featuring a slanted fascin crossband. (C) Schematic of a fascin crosslinked hexagonal bundle element. (D) Domain architecture of human fascin 1. (E) Composite 3.1 Å resolution cryo-EM density map of fascin crosslinked F-actin. (F) Atomic model of fascin crosslinked F-actin in ribbon representation.

## Results

### Structure of fascin bridging actin filaments

To gain insights into fascin’s F-actin binding and crosslinking mechanisms, we performed cryogenic electron microscopy (cryo-EM) studies of F-actin bundles generated by human fascin-1 (Methods). Bundles are not stoichiometrically defined protein complexes and do not possess helical symmetry, rendering them refractory to traditional filament image processing approaches. We therefore employed a neural-network–based particle picking strategy which detects positions within bundles featuring bridgeable pairs of filaments^58^, followed by asymmetric single-particle analysis (Methods, Table S1). We initially obtained a consensus reconstruction containing one fascin molecule crosslinking two parallel actin filaments at 3.4 Å resolution (Figure S1A). However, this density map featured streaking artifacts indicative of conformational heterogeneity. We therefore performed two rounds of multi-body refinement, alternately masking fascin together with each filament as one body while the other filament was considered a separate body. This procedure substantially improved the resolution of each actin-binding interface within the mask, yielding density maps at 3.0 Å and 3.1 Å resolution, while the other interface became blurred (Figures S1A-1C). A composite map was generated by aligning the fascin region of these two density maps, then combining the well-resolved interfaces while removing the blurred map regions. This composite map was used for atomic model building and refinement (Figure S1A).

The structure shows fascin employs two distinct ABS to crosslink a pair of actin filaments with an inter-filament distance of ∼12 nm. The overall shape of fascin is similar to that observed in the crystal structure of the protein in isolation^29–31^, with its four tandem β-trefoil domains arranged into a compact, bent horseshoe-shaped conformation (Figures 1D-1F). It consists of two pseudo two-fold symmetry-related lobes: one lobe comprising β-trefoils 1 and 2 and the other β-trefoils 3 and 4. The two distinct ABS are located on opposing surfaces of fascin at the interfaces between these lobes, with β-trefoils 1 and 4 forming ABS1, while β-trefoils 2 and 3 compose ABS2 (Figure 1F).

### Fascin undergoes rigid-body domain rearrangements to bind F-actin that are blocked by G2

When compared to the crystal structure of fascin in isolation (PDB: 3LLP, which we refer to as the “prebound” state)^29^, the conformation of each of the β-trefoil domains is nearly identical after F-actin binding (Figure S2A). However, substantial inter-domain rearrangements occur. The lobe composed of β-trefoils 1 and 2 rotates 23.6° towards the lobe composed of β-trefoils 3 and 4, facilitated by the flexible β-trefoil 2–β-trefoil 3 linker (residues 256-261). Additionally, β-trefoil 1 undergoes an independent 27.3° rotation via a hinge region (residues 137-141) connecting β-trefoils 1 and 2 (Figure 2A; Video S1). These inter-domain rotations substantially remodel fascin’s exposed surface, sculpting the two ABS into conformations competent for actin binding.

**Figure 2.**
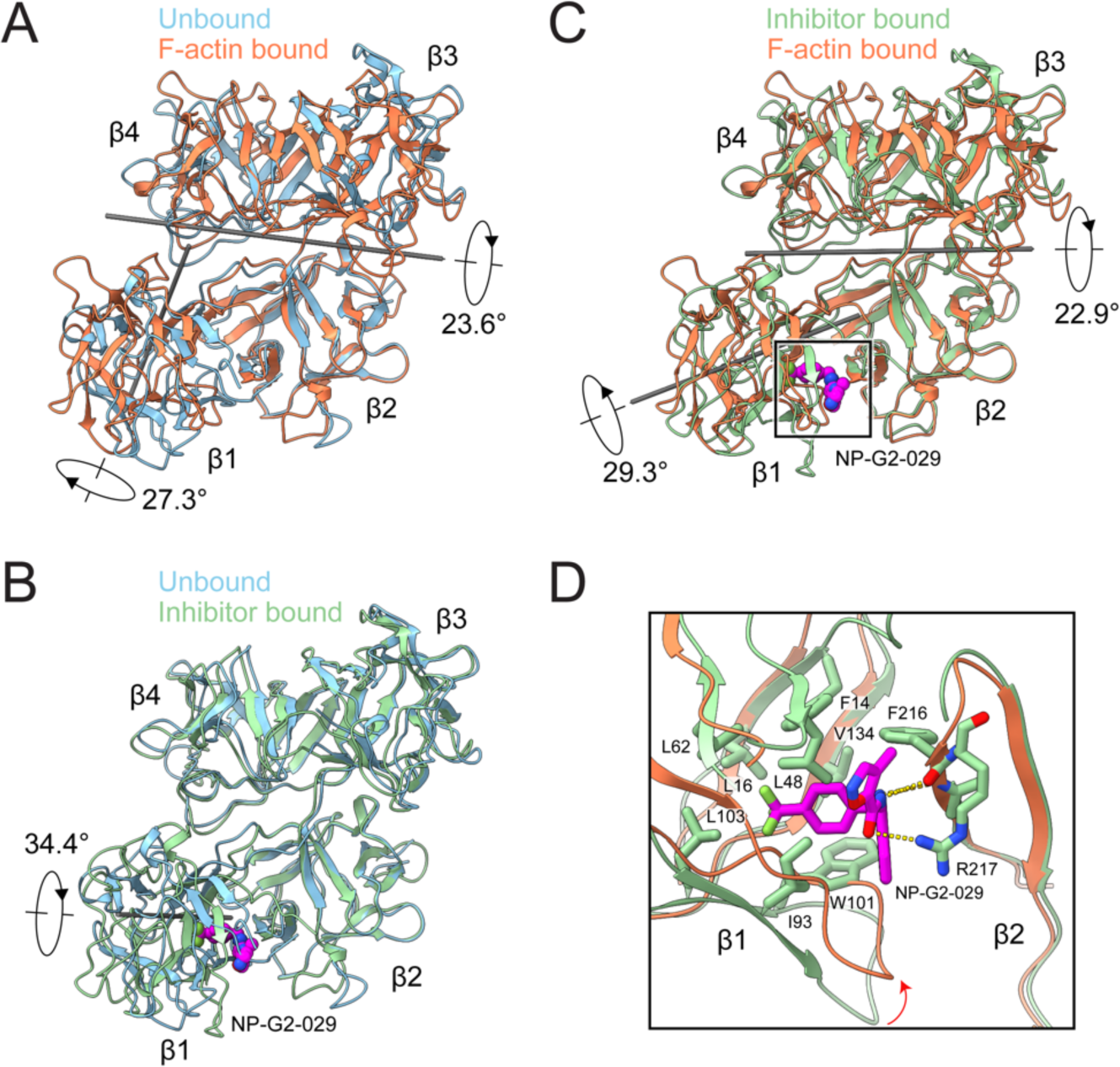
Fascin rearrangements upon F-actin binding are allosterically blocked by G2. (A) Superposition of fascin in the prebound (PDB: 3LLP) and F-actin bound states. (B) Superposition of fascin in the prebound and inhibitor-bound (PDB: 6B0T) states. The inhibitor NP-G2-029 is highlighted in space-filling representation. (C) Left: Superposition of fascin in the inhibitor bound and F-actin bound states. (D) Detail view of boxed region in C highlighting clash that would occur between NP-G2-029 and β-trefoil 1 in the F-actin bound state. All structures are aligned on β-trefoil 2.

We next sought to gain insight into the mechanisms of fascin inhibition by the G2 family of small molecules, which are currently undergoing phase 2 clinical trials for treatment of metastatic gynecological and breast cancers^43^. Previous structural studies of fascin bound to G2 in the absence of F-actin (PDB: 6B0T) showed that inhibitor binding induced a substantial 34.4° rotation of β-trefoil 1 via the same hinge region connecting β-trefoils 1 and 2^59^, suggesting that G2 might disrupt formation of the actin-binding conformation (Figure 2B; Video S1). Consistently, β-trefoil 1 undergoes rotations with distinct directions and magnitudes in the G2 bound structure and our F-actin bridging structure versus the prebound state. This results in its orientation differing between the actin-bound and G2 bound structures by 29.3°, disrupting the formation of ABS1 (Figures 2C and 2D; Video S1). As G2 binds in a cleft between β-trefoils 1 and 2 distal from both ABS, we infer that it acts as a molecular wedge, which allosterically blocks the inter-domain rearrangements required for fascin to adopt its actin-binding conformation.

### Fascin’s F-actin binding interfaces

Like many F-actin binding proteins, both fascin ABS engage a cleft formed by two longitudinally adjacent actin subunits along the same F-actin strand, here designated as subunits i and i+2 on F-actin 1 (which contacts ABS1) and subunits j and j+2 on F-actin 2 (which contacts ABS2; Figures 1F and 3A). This cleft is composed of each subunit’s subdomain 1, as well as the D-loop of subunit i+2 / j+2. At the fascin binding site, these two clefts on F-actin 1 and 2 are facing inwards towards the crossbridge. They are related by a shift of ∼27 Å (approximately F-actin’s helical rise) along the co-linear filament axes and a rotation of ∼ -167° (approximately F-actin’s helical twist). Fascin’s bent horseshoe architecture provides essentially perfect shape complementarity with this bi-partite, doubly spiraling interface, enabling each β-trefoil domain to form extensive contacts that maintain the two filaments in a parallel orientation with precise spacing. However, the two binding interfaces are highly asymmetric despite the overall shape complementarity between the two fascin ABS and the opposed F-actin clefts. Since F-actin 1 and F-actin 2 are superimposable, both of fascin’s ABS can engage the opposite filament by re-orientating the molecule, equivalent to the view shown in Figure 1F. Therefore, a fascin crossbridge has two fundamental binding poses, an “up” orientation (Figure 1F, left) and a “down” orientation (Figure 1F, right).

**Figure 3.**
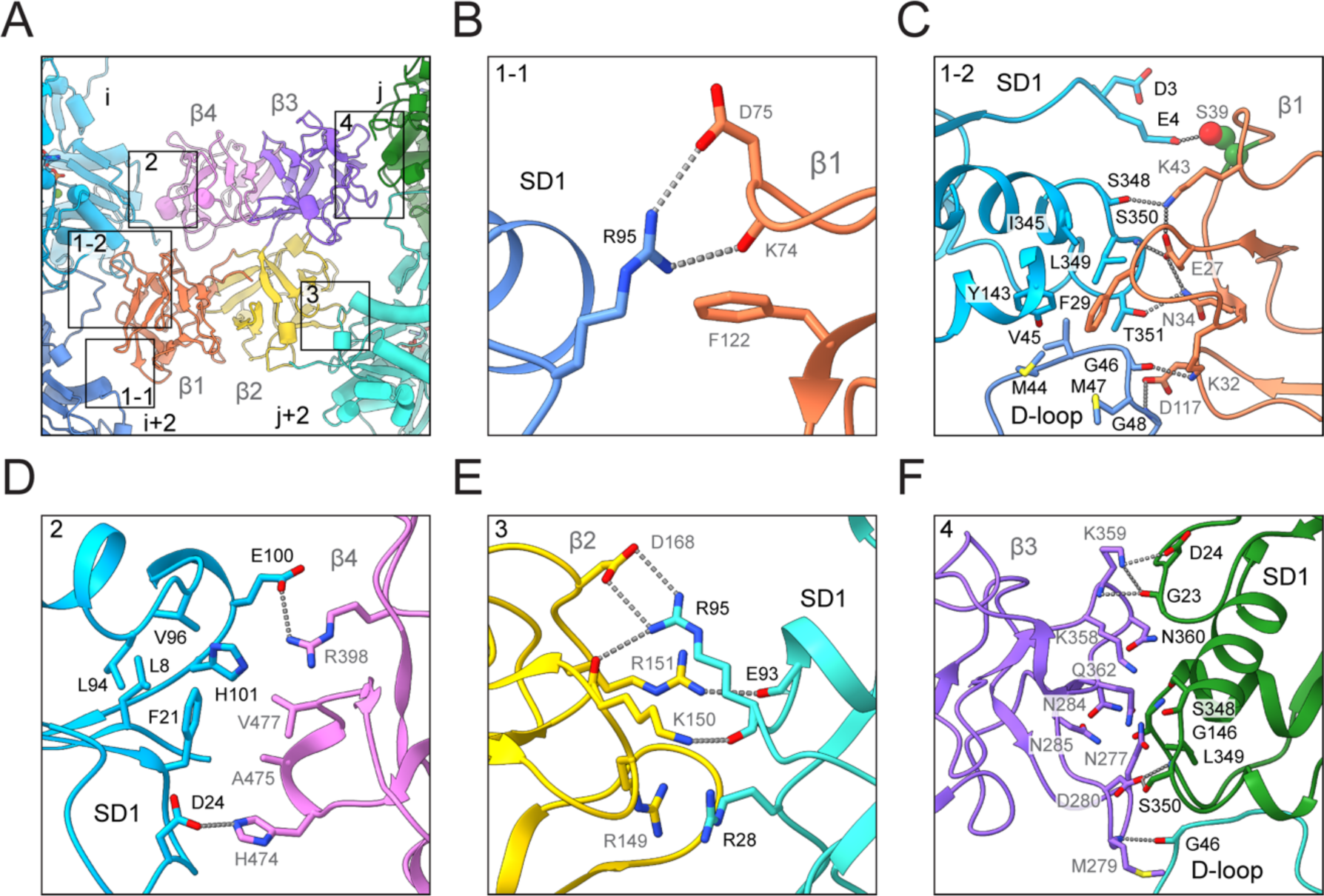
Fascin’s two F-actin binding interfaces are chemically distinct. (A) Overview of fascin’s two F-actin binding interfaces. (B) Contacts between subdomain 1 (SD1) of actin subunit i+2 and fascin β1. (C) Contacts between SD1 of actin subunit i, D-loop of subunit i+2, and fascin β1. PKC target residue S39 of fascin is highlighted in space-filling representation. (D) Contacts between SD1 of actin subunit i and fascin β4. (E) Contacts between SD1 of actin subunit j+2 and fascin β2, including cation clustering interaction between fascin R149 and actin R28. (F) Contacts between SD1 of actin subunit j, D-loop of subunit j+2, and fascin β3.

While all four of fascin’s structurally homologous β-trefoil domains are restricted to contacting actin subdomain 1 and the D-loop, the specific residue-level interactions they form are distinct (Figure 3A). β-trefoil 1 makes extensive contacts with two separate patches on F-actin 1. Towards the plus (“barbed”) end, fascin K74, D75 and F122 engage with a single actin residue, subunit i+2’s R95, through both hydrogen bonds and Van der Waals interactions (Figure 3B). K74 / D75 or F122 mutations were previously reported to have no effect on fascin’s bundling activity^32^, suggesting this interface is functionally nonessential. Towards the minus (“pointed”) end, β-trefoil 1 tightly packs against a large surface formed by subunit i+2’s D-loop and subunit i’s subdomain 1 through an extensive network of hydrophobic and electrostatic interactions (Figure 3C). Notably, the highly conserved fascin residue F29 is deeply buried in the hydrophobic groove at the inter-subunit longitudinal interface. Consistently, the F29A mutation severely impairs fascin’s bundling activity^32^. Fascin S39, the site of inhibitory PKC phosphorylation^41^, is adjacent to the N terminus of subunit i, where it forms a potential hydrogen bond with actin E4. Addition of a phosphoryl group on S39 would electrostatically repel the highly negatively charged actin N terminus, providing a mechanistic explanation for how this modification suppresses F-actin bundling. Several other fascin residues critical for F-actin bundling, including E27 and K43^32^, also mediate interactions at this interface. β-trefoil 4 exclusively interacts with subunit i’s subdomain 1, at a surface located above the actin N terminus (Figure 3D). Fascin residues A475 and V477 embed into a shallow hydrophobic groove on actin, while residues R398 and H474 form salt bridges with actin residues E100 and D24, respectively, consistent with R398 mutations abolishing fascin’s bundling activity^32^.

Like β-trefoil 4, β-trefoil 2 solely interacts with subdomain 1 of subunit j+2 on F-actin 2, but forms completely different contacts (Figure 3E). Fascin residues R151 and D168 form salt bridges with actin residues E93 and R95, respectively. R151 and D168 mutations have no effect on bundling activity^32^, indicating these interactions are auxiliary. Notably, fascin R149 and actin R28 form an electrostatics-defying *π*-cation / *π*-cation interaction through cation clustering^60^. Mutation of R149 severely impairs fascin bundling activity^32^, suggesting the unexpected interaction between this arginine pair plays a major role in mediating β-trefoil 2’s interaction with actin. Like β-trefoil 1, β-trefoil 3 makes extensive interactions with the interface between subunits j and j+2 on F-actin 2 (Figure 3F). However, the interface is dominated by extensive electrostatic interactions and Van der Waals contacts with subunit j’s subdomain 1, while the sole interaction with subunit j+2’s D-loop is a hydrogen bond between the backbones of fascin M279 and actin G46. In summary, we find that ABS1 and ABS2 form completely divergent contacts with F-actin despite fascin’s pseudo two-fold symmetry.

### Hexagonal bundle elements feature variable crosslinking architecture

We next sought to determine how fascin crosslinks helical F-actin filaments into hexagonal arrays despite their incompatible symmetries. We focused our analysis on “bundle elements”, minimal hexagonal sets of one central and six peripheral fascin-crosslinked filaments, from which higher-order arrays are built. After extensive classification, we recovered a single bundle element class spanning one F-actin crossover length (representing only 0.3% of the initial picks) which featured substantial fascin occupancy at all bridgeable positions. We refined this class to obtain an 8.7 Å reconstruction (Figures 4A, S2B and S2C). The overall organization of this bundle element is similar to those previously reported at lower resolution from tomographic studies, where the detailed architecture of the component filaments and crossbridges could not be determined reliably^32,46,54^. Within the bundle element, the seven filaments on the vertices of the hexagonal lattice result in twelve equally spaced filament pairs that are ∼12 nm apart, each of which is crosslinked by a single fascin. As anticipated, the twelve fascins are aligned along the three diagonals of the hexagon (Figures 4A and 4B). Each set of four fascin crossbridges oriented along the same diagonal (different colors in Figure 4A) occupy an approximate plane perpendicular to the aligned filament axes (a transversal layer), and the three transversal layers are roughly evenly spaced along the span of the crossover (Figures 4A and 4B). However, our subnanometer-resolution density map reveals that the four fascins comprising each layer are not perfectly aligned to form a linear crossband, instead exhibiting orientational and positional heterogeneity.

**Figure 4.**
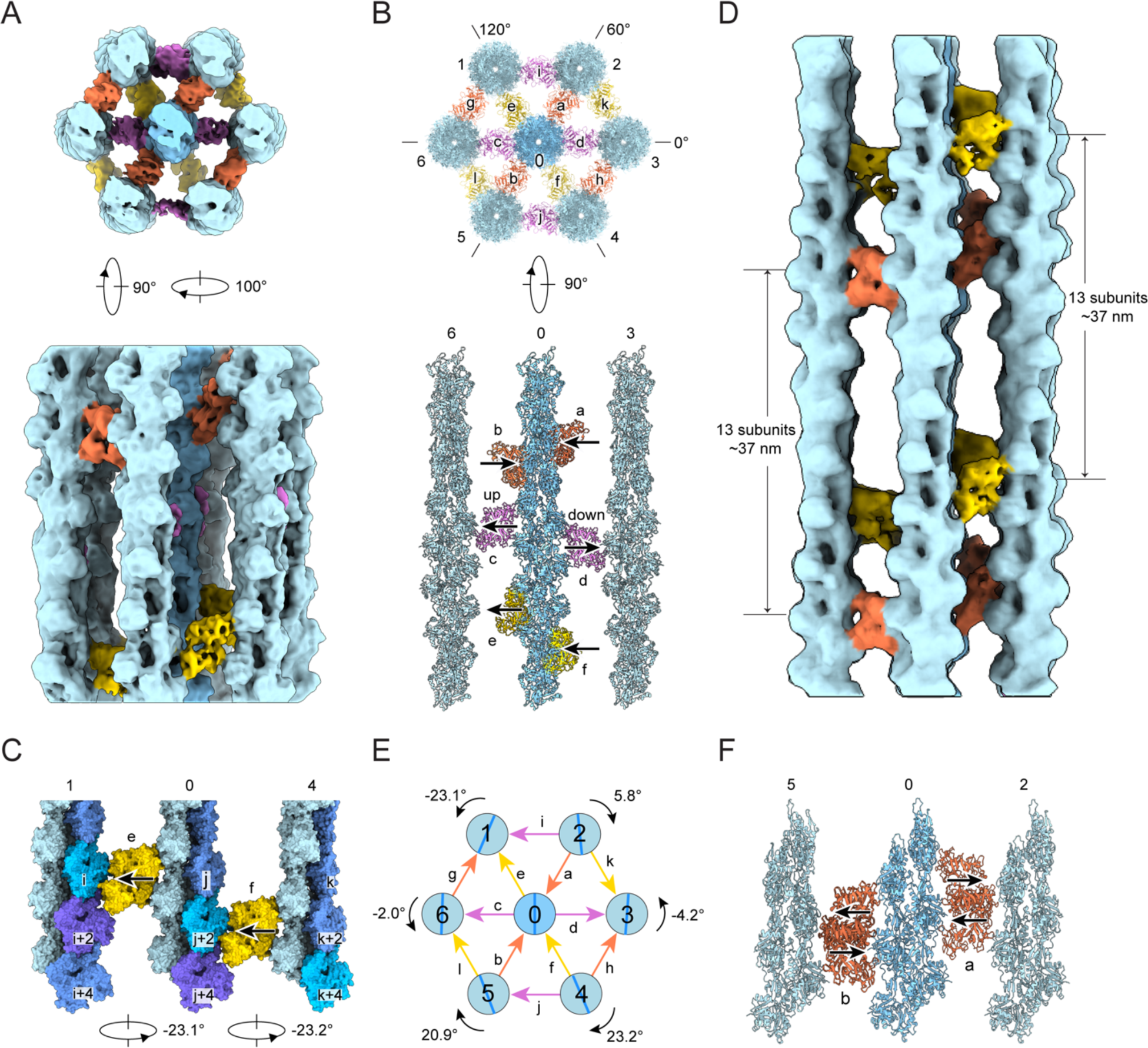
Architecture of a fascin crosslinked F-actin hexagonal bundle element. (A) End-on (top) and side view (bottom) of 8.7 Å resolution fascin crosslinked F-actin bundle element reconstruction (box size: 460 Å). Actin filaments are colored in shades of blue and fascins comprising each transversal layer are colored in different hues. (B) End-on (top) and side view (actin plane 6-0-3, bottom) of bundle element docking model in ribbon representation. The “up” and “down” poses of fascin c and d are indicated by arrows, with the arrowhead pointing towards ABS1. This orientation designation is used throughout the remainder of the manuscript. (C) Side view of actin plane 1-0-4, highlighting axial shift of fascin crossbridges associated with sequential filament rotational phase shifts. (D) Side view of 12.0 Å resolution bundle element reconstruction from particles reextracted with a box size of 740 Å. (E) Diagram of peripheral filament rotational phase shifts relative to the central filament, as well as fascin crossbridge poses. (F) Side view (actin plane 5-0-2) of five bundle element docking models superimposed on their central filaments, highlighting variations in fascin pose and axial positioning at equivalent bridging locations.

Each transversal layer features fascin crossbridges in both the “up” and “down” poses implied by our high-resolution crossbridge structure (Figure 1F), with the two modes distributed throughout the bundle element without discernable regularity (Figures 4B, S2E and S2F; Video S2). For example, in the actin plane spanning filaments 6-0-3, fascin c is “up”, while fascin d is “down” (Figure 4B, bottom panel). Additionally, within a transversal layer, fascins featuring the same binding mode can be displaced along the longitudinal axis. For example, in the actin plane spanning filaments 1-0-4, the two “up” oriented fascins e and f are longitudinally offset by one actin subunit (Figures 4C and S2E). The combination of the two fascin crosslinking poses and shifts in fascin axial positioning gives rise to the noncoherent arrangement of crossbridges across transversal layers. While our bundle element reconstruction reveals substantial lateral disorder, the helicity of F-actin could nevertheless give rise to longitudinally repetitive binding patterns. To probe this potential periodicity, segments were re-extracted with a larger box size of 720 Å, spanning approximately two F-actin crossovers, and refined to yield a 12.0 Å reconstruction (Figures S2B and S2D). The pose and position of each fascin precisely repeats in the next crossover (Figure 4D; Video S2), indicating a longitudinal periodicity of ∼37 nm, consistent with previous *in vitro* and *in situ* studies^31,32,46,54,55^. However, variability in F-actin’s helical twist may erode this longitudinal order at longer length scales^61,62^. Collectively, our data suggest flexibility in fascin’s binding pose and axial positioning enables the protein to accommodate a range of inter-filament geometries encountered at different bridging positions.

### Fascin crosslinked filaments are rotationally noncoherent

Early theory predicted the emergence of linear crossbands when bundled actin filaments are in perfect register^46,52^. As we instead observe only partially ordered transversal layers featuring axially shifted fascins, we hypothesized that the filaments in our bundle element reconstruction are rotationally noncoherent. We therefore measured the rotational phase shift of the six peripheral filaments in relation to the central filament, which we define as the axial rotation required to superimpose a filament with the central reference when viewed from the minus end. We found that all six peripheral filaments exhibit distinct rotational phase shifts that vary in direction and magnitude, which can be broadly classified into two groups (Figure 4E).

One group, consisting of filaments 2, 3, and 6, displays minor phase shifts of 5.8°, -4.1° and -2° respectively. As these values approximate 0°, the filaments in the 6-0-3 actin plane are nearly in register, with highly similar binding interfaces between filament pairs 6-0 and 0-3 (Figure 4B, bottom panel). However, fascin c, crosslinking filament pair 6-0, adopts the “up” pose, while fascin d, crosslinking filament pair 0-3, adopts the “down” pose. This suggests that pairs of filaments which are rotationally in-phase can nevertheless be crosslinked by either binding mode, a degeneracy anticipated to produce crossband disorder even in well-registered filament arrays. The other group includes filaments 1, 4, and 5, which display substantial rotational phase shifts of -23.1°, 23.2° and 20.9°, respectively. Given the helical symmetry of F-actin, a rotation of ±26.7° is equivalent to a translation of one full actin subunit up or down the filament axis. In the 1-0-4 actin plane, two such consecutive rotations produce equivalent fascin crosslinking sites axially shifted by one actin subunit between the 1-0 and 0-4 filament pairs. Fascins e and f bind at these sites, both in the “up” pose, resulting in a prototype of a slanted crossband (Figures 4C and S2E).

As fascin’s crosslinking geometry appears to vary with rotational phase shifts between filaments, we hypothesized that other bundle elements with alternative inter-filament rotation / crosslinking patterns could be present in our data. We therefore selected four additional bundle element classes for refinement (Figure S2B). While these reconstructions varied in quality, they were all sufficiently detailed to reliably model the central actin filament and its six fascin crossbridges. We then superimposed these four bundle element models on the central filament of the initial bundle element model. All five models feature fascin crossbridges clustered at six distinct locations consistent with three transversal layers (Figures S2G-S2I). At each location, both the “up” and “down” crosslinking poses occur with varying frequencies (Figures 4F and S2G-S2I). None of the classes share the same distribution of fascin poses, suggesting each class represents a unique bundle element geometry (Figures 4E and S2J). We furthermore observed that for a given transversal layer, different classes featured pairs of fascins with the same pose that were either co-linear or axially shifted by one actin subunit, suggesting varying rotational phases between their component filaments. Thus, we infer that fascin’s two binding poses coupled with filament rotational freedom enable the assembly of hexagonal bundle elements with varying crosslinking geometries. Moreover, the occupancy of individual fascins and filaments varies substantially between bundle element classes (Figure S2B), implying differential stability.

### Fascin structural plasticity facilitates flexible crosslinking

Beyond the two binding poses we identified, fascin’s capacity to accommodate geometric variations between binding positions has been predicted to require crossbridge structural plasticity^52^. We therefore performed detailed analysis of the multibody refinement that led to our high-resolution crossbridge structure. The major variability mode features a continuous counterclockwise rotation of F-actin 2 relative to F-actin 1 when viewed from the minus end, while their filament axes remain parallel. To assess the structural underpinnings of this variation, particles were partitioned into three distinct groups along the corresponding eigenvector, where their amplitudes are well-described by a Gaussian distribution (Figure 5A). Subsequent masked refinements resulted in three reconstructions at resolutions of 3.9 Å, 3.4 Å, and 4.0 Å, representing structural snapshots along the conformational heterogeneity continuum that we refer to as eigen_left, eigen_middle, and eigen_right (Figures 5B and S3A-S3C).

**Figure 5.**
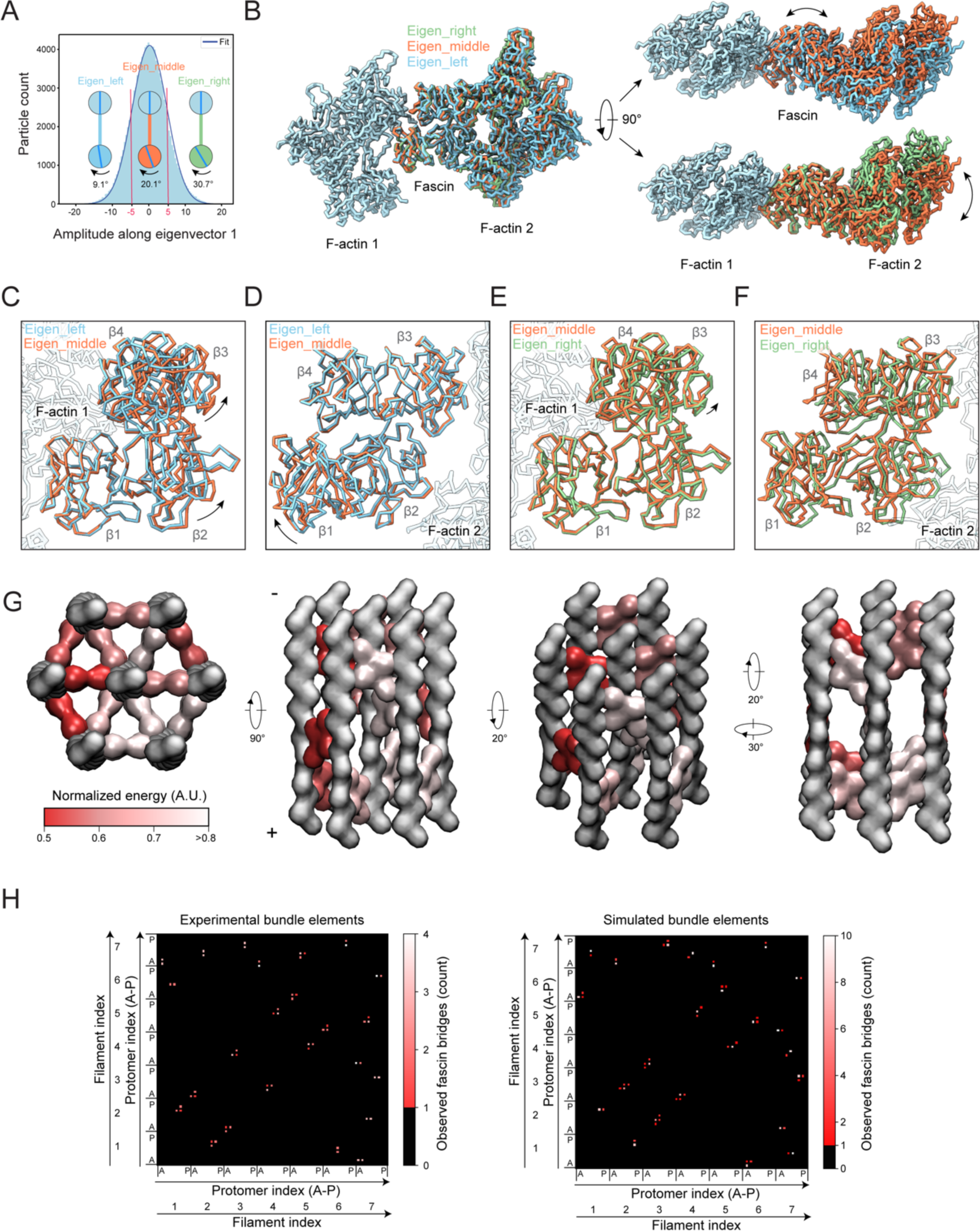
Fascin crossbridge structural flexibility mediates inter-filament rotations. (A) Histogram of amplitudes along the first multibody refinement eigenvector for all particles (n = 113,800), which follows a unimodal Gaussian distribution (R^2^ = 0.9986). Red lines indicate cutoffs for partitioning particles into three bins. Rotational phase differences between filaments from each reconstructed bin are depicted. (B) Side (left) and end-on (right) views of atomic models from eigenvalue binned reconstructions superimposed on F-actin 1. Predominant structural transitions explaining the relative rotation of F-actin 2 are indicated by double-headed arrows. (C and D) Structural comparison of eigen_left and eigen_middle atomic models when superimposed either on F-actin 1 (C) or F-actin 2 (D). Rigid body repositioning of fascin subdomains are indicated by black arrows. (E and F) Structural comparison of eigen_middle and eigen_right reconstructions when superimposed either on F-actin 1 (E) or F-actin 2 (F). (G) Views of a representative simulated hexagonal bundle element. Actin is shown in grey, and fascins are colored by normalized energy scores. (H) Contact maps of experimental (n = 5) and simulated (n = 10) hexagonal bundle elements. Models are aligned on the central filament. Each filament contains 16 protomers (A-P) indexed alphabetically from the barbed end to the pointed end. The peripheral filaments are numbered sequentially from 1-6, while the central filament is assigned index 7. Each matrix element depicts a potential fascin crosslinker position in the bundle, while color corresponds to the observed number of fascin crossbridges.

Comparison of atomic models refined into these maps reveals indistinguishable conformations for F-actin 1 and F-actin 2 across all three snapshots, indicating that F-actin structural plasticity does not substantially contribute to crossbridge flexibility (Figure S3D). Comparing the eigen_left and eigen_middle snapshots show no distinguishable changes at either of the ABS-actin interfaces, suggesting rearrangements within the fascin molecule itself are primarily responsible for this aspect of the conformational landscape (Figures 5C, 5D, S3E-K). Consistently, in the reference frame of F-actin 1, β-trefoil 1 remains stationary, while β-trefoils 2-4 undergo a rigid-body rotation through two pivot points: the F-actin 1–β-trefoil 4 interface and the flexible linker connecting β-trefoils 1 and 2 (residues 137-141), resulting in a 3 Å and 7 Å displacement of ABS2 and F-actin 2, respectively (Figure 5C; Video S3). Notably, the actin-binding residues on β-trefoil 4 are clustered in two pliant surface loops (Figures 3D and S3H), allowing β-trefoil 4 to rotate while maintaining interactions with the filament.

Conversely, when the eigen_middle and eigen_right snapshots are compared in the reference frame of F-actin 1, fascin’s conformation remains essentially unchanged (with only modest repositioning of β-trefoils 2-4 relative to β-trefoil 1), as does the ABS1–F-actin 1 interface (Figures 5B, 5E and S3E-H). However, alignment on F-actin 2 reveals substantial remodeling of the ABS2–F-actin 2 interface, with both β-trefoils 2 and 3 displaced relative to the filament surface (Figure 5F). At the β-trefoil 2–actin interface, key interactions are maintained by repositioning the side chains of residues located on flexible loops (Figure S3L). Notably, the fascin R149-actin R28 *π*-cation / *π*-cation interaction is preserved, suggesting this unusual contact may contribute to the interface’s plasticity. However, at the actin–β-trefoil 3 interface, both actin-interacting loops on β-trefoil 3 are displaced, suggesting this contact is destabilized in the eigen_right snapshot (Figure S3M). Taken together, this analysis suggests remodeling of both fascin itself and the ABS2–F-actin interface mediate crossbridge structural plasticity (Video S3).

We further examined the range of inter-filament rotations spanned by the three structural snapshots, which reflect the distribution of rotational phase shift magnitudes between filaments in the bundle elements from which the particles were extracted. With F-actin 1 as the reference, the eigen_left, eigen_middle, and eigen_right snapshots feature offsets of 9.1°, 20.1° and 30.7°, respectively (Figure 5A), suggesting the presence of substantial rotational heterogeneity in fascin crosslinked F-actin bundles. As the eigen_middle reconstruction comprises particles centered around the peak of the Gaussian distribution, this analysis indicates that rotational phase shifts with a magnitude of approximately 20° are the most probable (and by extension the most energetically favorable) for fascin crosslinked filament pairs. Our analysis of bundle elements found such rotations are associated with axial shifts in fascin positioning along transversal layers (Figure 4C), whose apparent favorability could mediate the formation of slanted crossbands in higher-order bundles. Nevertheless, fascin can clearly tolerate a broad range of interface geometries, albeit with reduced probability (and by extension with an incurred energetic cost), to circumvent symmetry mismatches while constructing hexagonal arrays from helical F-actin.

### A minimal model recapitulates variable bundle element architecture

To assess whether the notable structural features we identified are the major determinants of fascin bundle construction, we developed a minimal computational model. In our model, actin filaments are sequentially added to a bundle with rotational phase shifts that maximize the probability of fascin crosslinking. The probability of fascin binding at each potentially bridgeable pair of protomers between a newly added filament and pre-existing filaments is assessed through a scoring function parameterized by our multi-body refinement analysis. Protomers with scores above an empirical cutoff are initially nominated as candidates for fascin binding positions. From these candidate positions, the rotational shift corresponding to the maximum overall score is selected, and the corresponding protomers are assigned as having fascin bound (Methods, Figure S4).

We generated seven-filaments assemblies corresponding to bundle elements while varying the absolute rotational shift of the central filament (which cannot be assigned in our experimental bundle element reconstructions). We focused on assemblies featuring central filaments with absolute rotational shifts between 0-10° for detailed analysis. Consistent with our experimental reconstructions, the simulated assemblies feature substantial variability in the rotational phases of their component filaments and in the positioning of individual fascins, while nevertheless maintaining overall transversal layer organization (Figure 5G). This is visualized in an averaged contact map (Figure 5H), which consists of sparse peaks corresponding to allowable bridging locations that feature broadening due to variability in axial positioning. A contact map calculated from the experimentally observed bundle elements is similar, particularly for crossbridges contacting the central filament (Figure 5H). Discrepancies in outer filament crossbridging patterns are likely due to the reductionist nature of the model, as well as potential non-representative sampling by our 3D classification procedure, which only recovers classes which can be readily averaged. Nevertheless, the general correspondence between the model and our experimental data suggests that the structural features embedded in the model parameters, which encode fascin binding probability modulated by inter-filament rotation angles, can recapitulate the flexible three-dimensional arrangement of hexagonal bundle elements.

### Filament rotational phase shifts establish emergent fascin binding patterns

We next sought to visualize how the structural plasticity of bundle elements gives rise to emergent fascin binding patterns in higher-order bundle assemblies by analyzing *in vitro* reconstituted fascin-crosslinked F-actin bundles using cryo-electron tomography (cryo-ET). While cryo-ET theoretically has the potential to produce molecular-resolution three-dimensional reconstructions of micron-sized fields of view, this imaging modality has been practically limited by low signal-to-noise and non-isotropic resolution, with notable blurring in the z-dimension. To overcome these limitations, we substantially adapted our neural-network–based processing framework to denoise our tomograms and semantically segment fascin and F-actin (Methods, Figure S5). The unprecedented molecular details revealed by this procedure enabled us to unambiguously distinguish the three-dimensional positions and orientations of actin filaments and individual 50 kDa fascin molecules without averaging (Figure 6A; Video S4), facilitating direct analysis of bundle network geometry.

**Figure 6.**
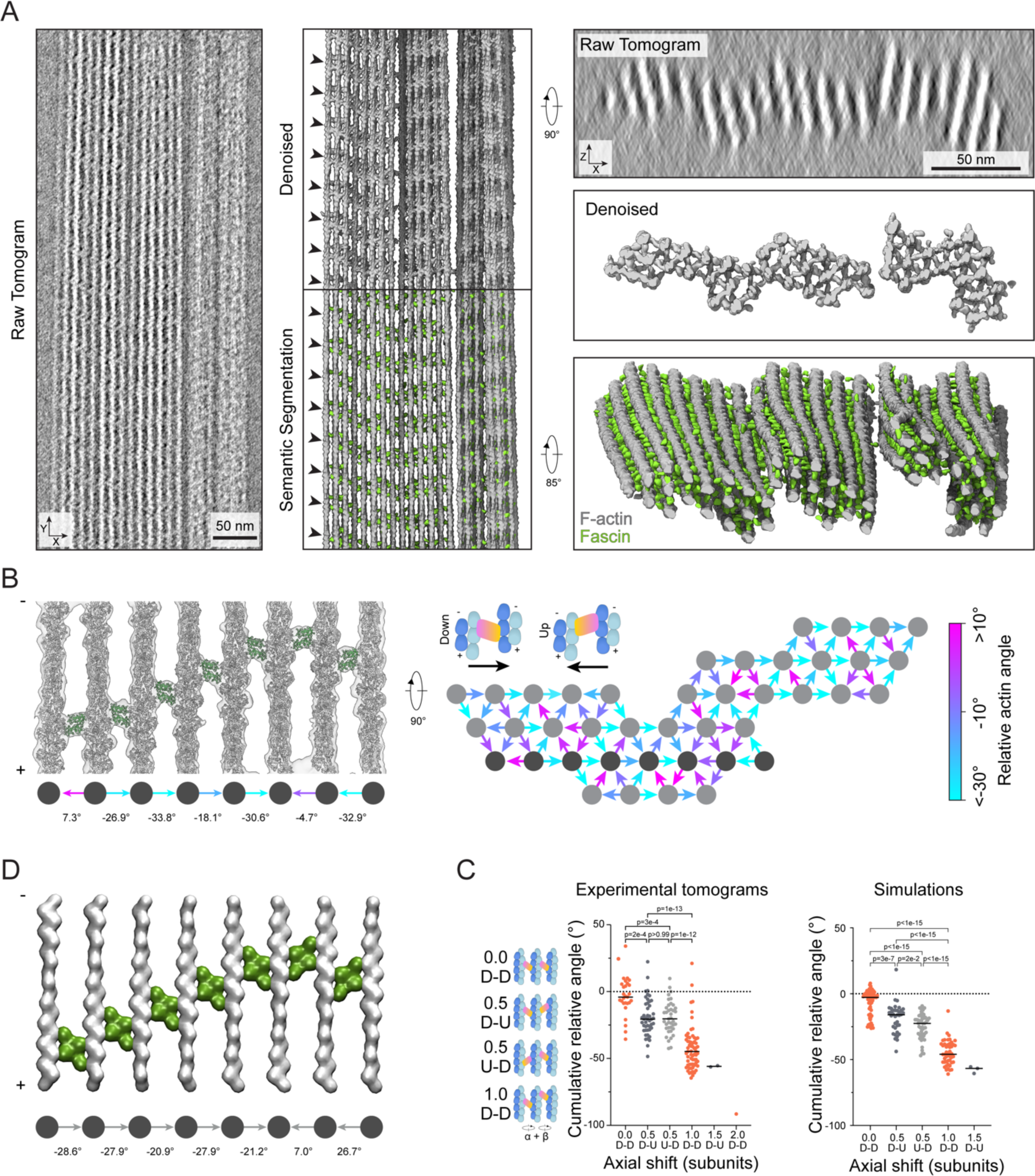
Denoised tomograms reveal nanoscale organization of fascin crosslinked bundles. (A) Left: Side views of projected raw, denoised (middle top) and semantically segmented (middle bottom) tomogram. Arrowheads indicate periodic chevron-like pattern of fascin crossbridges. Right: end-on views of projected raw (top) and denoised (middle) tomogram, as well as oblique view of semantic segmentation (bottom). (B) Left: Side view of representative rigid-body docking model of one actin plane from a denoised tomogram. Filament rotational phase shifts and fascin poses (arrows) are indicated. Right: End-on view schematic of filament rotational phase shifts and fascin poses throughout a 17-protomer thick slab of a bundle. Dark grey filaments correspond to actin plane displayed on the left. (C) Quantification of cumulative filament rotational phase shifts versus fascin axial shifts across observed three-filament configurations from denoised tomograms (n = 293 from N = 5 bundles) and simulations (n = 202 from N = 8 bundles). Orange points indicate matched fascin poses, while points in shades of grey indicate mismatched fascin poses. (D) Side view of the central actin plane from a representative simulation of a 22-filament bundle. Filament rotational phase shifts and fascin poses (arrows) are indicated.

In the denoised tomograms, we observed tightly packed parallel bundles that typically spanned the entire diameter of 1.2 μm holes in the carbon support film (Figures 6A, S6A, S6B). Bundles displayed nearly complete fascin decoration, with the ∼37 nm axial periodicity of fascin crossbands persisting for the entire region visualized in a typical straight bundle, spanning tens of crossovers (Figure 6A). They feature an oblong aspect ratio when viewed end-on; typically bundles were three to five layers thick along the z direction but could be over ten layers wide. Many bundles exhibit substantial morphological flexibility and spatial variations in filament composition at the micron scale. Frequently, a subset of filaments will splay from one bundle to become an independent bundle or twist and coalesce with other bundles (Figure S6A), similar to what has been observed in filopodia^54,55^. Moreover, in some bundles, all filaments collectively twist along the longitudinal axis (Figure S6B). Furthermore, multiple sparsely interconnected smaller bundles with distinct crosslinking patterns can coalesce into larger bundles (Figure S6A, S6B).

To study the relationship between bundle architecture and the organization of their molecular constituents, we selected four regular bundles composed of 29 to 41 straight filaments for detailed analysis. We were able to rigid-body dock atomic models of fascin crossbridges and 17-protomer actin filaments (spanning one crossover length) directly into the denoised tomograms, allowing us to assign fascin positions and poses while measuring corresponding filament rotational phase shifts (Figures 6B, S6C, S6D). We occasionally observed actin planes in which all filaments are consecutively rotated by approximately -25° (Figure S6D). Correspondingly, the fascins are all in the “down” pose and sequentially axially shifted to form a slanted crossband. More frequently, crossbands adopt a chevron morphology through a combination of consecutive approximately -25° rotations followed by a series of approximately +25° rotations (Figure 6B). Despite these features of short-range order, globally we observe extensive heterogeneity in the architecture of hexagonal bundle elements (Figures 6B, S6C) confirming our 3D classification analysis (Figures 4E, S2B, S2J).

We next sought to determine how these fascin binding patterns are propagated across bundles. We focused our analysis on co-planar sets of three actin filaments featuring two fascin crossbridges, minimal units which connect adjacent, partially overlapping hexagonal bundle elements. After standardizing the viewing orientation such that only unique views were retained, we identified a set of rules relating the cumulative rotational phase shift across the three filaments to patterns of fascin binding poses and axial offsets, which collectively produce chevron-shaped crossbands (Figure 6C). If the cumulative rotational phase shift is approximately 0°, both fascins will be in the same orientation and have no axial shift. If the cumulative rotational phase shift is approximately -25°, the fascins will be oppositely oriented and have an axial shift of half an actin subunit length, with no preference in the pose of the first fascin. If the cumulative rotational phase shift is doubled to approximately -50°, the fascins will be axially offset by the length of one full actin subunit and both be in the same (down) pose. As these cumulative rotational phase shift values are nearly multiples of the rotation linked to a one subunit shift along a single filament (±26.7°), these patterns likely emerge to accommodate both F-actin’s helical symmetry and plausible crosslinking positions in the hexagonal bundle lattice. Rarely, more extreme shifts were observed, with axial shifts of 1.5 or 2 subunit lengths, which continued the trend of oppositely oriented fascins for half-subunit length shifts and equivalently oriented fascins for full subunit length shifts.

Notably, there is some overlap in the cumulative phase shifts between configurations featuring distinct fascin axial positionings (Figure 6C), consistent with fascin’s capacity to adopt either the up or down binding pose when the magnitude of the rotational phase shift between two filaments is low. This orientational flexibility introduces wobble positions in the system that facilitate the formation of chevrons by switching the direction of slanted crossbands. This likely contributes to overall bundle stability by allowing fascin to tolerate diverse binding site geometries and corresponding inter-filament architectures.

We next assessed whether similar binding patterns can emerge from our minimal model by simulating bundles of 22 filaments arranged into three layers (Methods). Indeed, we found substantial enrichment of slanted and chevron-shaped crossbands, visible in both individual assemblies (Figure 6D), and in the positioning of fascins across trials, which reproduced our experimentally determined binding patterns (Figure 6C). This suggests geometric linkages between binding sites produce long-range binding patterns, coordinating the local specification of fascin binding positions and poses across higher-order assemblies.

### The accumulation of unfavorable inter-filament interfaces restricts bundle size

We next sought to gain insights into mechanisms which limit the size of fascin-cro sslinked bundles, which generally do not exceed 40 filaments^45,49,54,55^. We reasoned the geometric diversity we observed in our tomograms would also give rise to a range of crossbridge conformations which vary in their frequency and corresponding energetic favorability. To survey the crossbridge conformational landscape in our cryo-ET data, we pursued subtomogram averaging and variability analysis. Our semantic segmentation results facilitated automated extraction of subtomograms well-centered on fascin molecules (Figure S7). Subtomogram averaging produced a 6.7 Å consensus reconstruction of two filaments cross-linked by fascin (Figures S7A, S7B, S7D, Table S2) with 3D classification indicating minimal false-positive picks, validating our denoising approach (Methods). Subsequent multi-body refinement and flexibility analysis revealed highly similar deformation modes to those we observed in our single particle analysis. Specifically, hinging motions dominated variability, while purely translational displacements were minimal (Figures S7C, S7E). The deviation of each subtomogram from the consensus structure was measured using the Mahalanobis distance, a metric which allowed us to simultaneously consider all eigenvectors corresponding to rotational displacements (Methods, Figure S7F). We then mapped these scores back to the corresponding particle positions in the full tomograms to generate spatial maps of the relative favorability of individual fascin crossbridges within bundles (Figure 7A).

**Figure 7.**
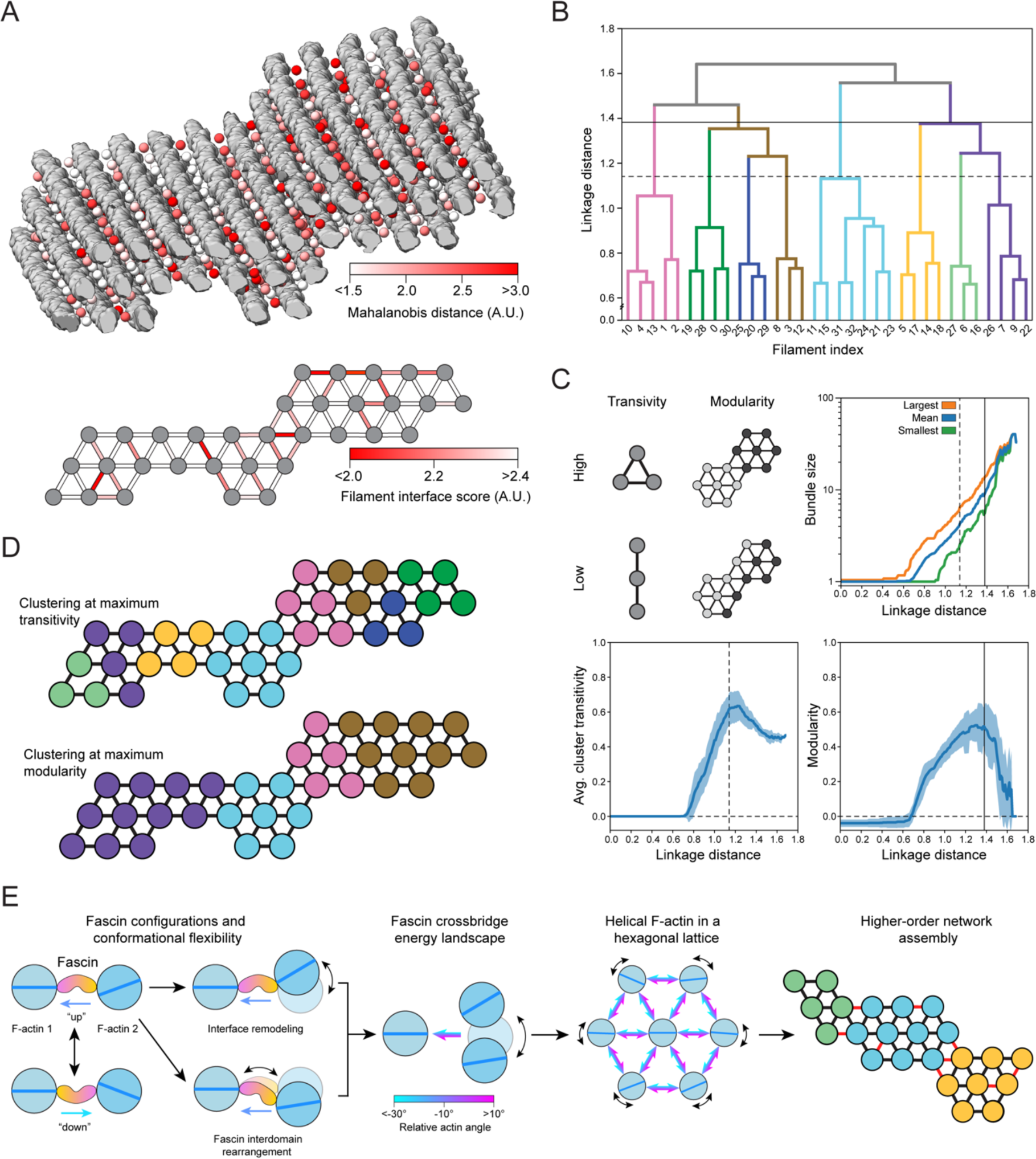
Accumulation of unfavorable crossbridges restricts bundle growth. (A) Top: Oblique view of fascin crossbridge Mahalanobis distances calculated from multibody refinement (colored spheres) mapped on to F-actin bundle. Filaments are displayed in grey. Bottom: Bundle filament interface scores represented as a weighted graph. (B) Dendrogram of graph-based hierarchical clustering of the bundle represented in A; colors represent clusters at linkage distance corresponding to maximum transitivity. (C) Plots of graph metrics versus linkage distance. Top right: Quantification of largest, mean, and smallest average cluster sizes across bundles. Bottom: Average cluster transitivity (left) and modularity (right); error bars indicate s.d. Vertical dashed and solid lines indicate linkage distances with maximum transitivity and modularity, respectively, for the bundle shown in A. N = 24 bundles. (D) Graphical representation of the clustering at maximum transitivity (top) and modularity (bottom) for the bundle shown in A. (E) Cartoon detailing the link between nanoscale conformational flexibility of the fascin crossbridge and variable micron-scale bundle architecture, which limits bundle size.

Inspection of these maps did not reveal a clear pattern at the level of individual fascins, consistent with our observation of extensive geometric heterogeneity across bundles (Figure S6A, S6B, S6C). As the overall cohesion between filaments in a bundle is mediated by the cumulative action of the crosslinkers at each inter-filament interface, we examined variations at this intermediate scale. To simplify the analysis, we considered 24 discrete bundle regions of uniform length. The interquartile range for the number of filaments per bundle was 23 to 43 filaments. We reduced each region to a graph representation consisting of filament nodes and interface edges. An interface score was then calculated for each edge, which is a function of the number of crossbridges along that interface and their average Mahalanobis distance (Methods, Figure 7A). This revealed variations in interface favorability which were irregularly distributed across bundles.

We then performed hierarchical clustering of the graphs to investigate how nonuniform interface stability impacts bundle organization (Figure 7B), a procedure which sequentially divides each bundle into smaller clusters through preferential partitioning across unfavorable interfaces. We analyzed the resultant dendrograms through the metrics of average cluster transitivity and modularity, which serve as proxies for cluster stability that are independent of cluster size above a minimum threshold of three filaments (Figure 7C). Transitivity reflects the number of connected triangles, which form the basis for actin’s hexagonal tiling, while modularity reflects the strength of connections within versus between clusters. When analyzed across bundles, both metrics feature a distinct peak at intermediate linkage distance, suggesting optimum stability of clusters smaller than full bundles. These range in average size from four (transitivity peak) to eight (modularity peak) filaments, with a minimum of three (transitivity peak) and a maximum of 15 (modularity peak; Figures 7C, 7D). Inspection of the connection patterns revealed that the boundaries between clusters at the modularity peak corresponded to the worst interfaces (Figure 7D), supporting a model in which these represent weak points within bundles. This suggests that lateral expansion of bundles results in the accumulation of unfavorable interfaces, with larger bundles being composed of irregular mosaics of stable clusters. While this analysis does not provide a quantitative explanation for the precise size distribution of fascin crosslinked bundles, it does suggest there is an energetic barrier to their indefinite expansion, an intrinsic size restriction mechanism imposed by their variable molecular architecture.

## Discussion

This study integrates atomistic structural characterization of the flexible fascin–F-actin interface with network-level analysis of bundle architecture, allowing us to uncover multi-scale mechanistic principles for the construction of a supramolecular cytoskeletal assembly (Figure 7E). Our structural analysis provides detailed insights into conformational dynamics underlying fascin’s regulated F-actin engagement and their allosteric inhibition by G2, providing a foundation for guiding the development of optimized fascin inhibitors. At the network scale, we find that fascin’s two F-actin binding poses and crossbridge conformational flexibility facilitate the incorporation of helical actin filaments into a hexagonal bundle lattice by accommodating a broad range of inter-filament rotation angles. This insight was enabled by the cryo-ET denoising procedure we introduce here, which allowed us to establish explicit links between protein-level structural plasticity and bundle network geometry. Our denoising approach has limitations, notably the requirement for prior structural knowledge of all system components, and thus in its current form it is primarily applicable to reconstituted preparations. Nevertheless, our work highlights the potential for structural dissection of large macromolecular ensembles composed of hundreds to thousands of proteins, where functional conformational dynamics can emerge that would remain inscrutable in isolated components.

Our data further shed light on how flexible fascin–F-actin network assembly gives rise to long-range binding patterns and restricts bundle size through the introduction of unfavorable interfaces, emergent phenomena which are both mediated by the links between fascin’s conformational landscape and variable bundle architecture. While here we focused on ground-state network geometry, filopodia primarily function in mechanically active environments. Our data imply that forces which deform fascin crosslinked bundles could alter their internal molecular structure and functional properties, for instance by rearranging the F-actin helix through twist-bend coupling^63,64^, as well as by modulating fascin’s conformational landscape. Additionally, while we find that the minimal two-component fascin / F-actin system recapitulates much of the previously described complexity of filopodia cores^8,11^, fascin collaborates with additional crosslinkers to build other classes of protrusions such as stereocilia^65,66^. One such crosslinker is plastin, whose conformational dynamics differ substantially from fascin’s^7,58,67^, suggesting multi-crosslinker networks may feature distinct architectures and functional properties. Indeed, in the presence of fascin, the addition of another crosslinker allows bundles to grow to larger sizes through unclear mechanisms^49^. We anticipate the multi-scale structural approach we introduce here will enable future work examining multi-component cytoskeletal networks, allowing detailed examination of how their nanoscale conformational dynamics intersect with their mesoscale architecture, mechanical properties, and cellular functions.

## Acknowledgements

We gratefully acknowledge Johanna Sotiris, Honkit Ng, and Mark Ebrahim at the Rockefeller University Evelyn Gruss Lipper Cryo-EM Resource Center (CEMRC) for their assistance with data collection, and Ariel Pan (RU) for performing F-actin purification and assistance with preparing an interface for neural-network based particle picking. We thank Lynda Chin for providing the cDNA of human fascin 1 via Addgene (plasmid # 31207). This work was funded by grants from the NIH (R01GM141044), the Alfred P. Sloan Foundation (G-2020-14047), and the Stavros Niarchos Foundation Institute for Global Infectious Disease Research at the Rockefeller University to G.M.A. as well as NIH grant R35GM147481 to T.C.B.

## Resource Availability

All resources and reagents reported in this study are freely available, and requests should be directed to the lead contact, Gregory M. Alushin.

## Data Availability

The cryo-EM density maps and atomic models generated in this study have been deposited in the Electron Microscopy Data Bank (EMDB) and Protein Data Bank (PDB): fascin bound filament 1 (EMDB: EMD-43364; PDB: 8VO5); fascin bound filament 2 (EMDB: EMD-43365; PDB: 8VO6); composite map of the fascin crossbridge (EMDB: EMD-43366; PDB: 8VO7); multibody binned reconstructions eigen_left: (EMDB: EMD-43367; PDB: 8VO8); eigen_middle: (EMDB: EMD-43368; PDB: 8VO9); eigen_right: (EMDB: EMD-43369; PDB: 8VOA); fascin crosslinked hexagonal bundle element with a box size of 460 Å (EMDB: EMD-43370); fascin crosslinked hexagonal bundle element with a box size of 740 Å (EMDB: EMD-43371); subtomogram average of fascin crossbridge (EMDB: EMD-43372). The trained neural networks for denoising and semantically segmenting micrographs and tomograms as well as the atomic models and volume maps used to generate synthetic datasets are available at Zenodo: https://doi.org/10.5281/zenodo.10456803. All other data are presented in the manuscript.

## Code Availability

Custom code is available at https://www.github.com/alushinlab/fascin as open source.

## Author contributions

R.G., M.J.R, T.C.B., and G.M.A. designed research. R.G. purified fascin and collected cryo-EM and cryo-ET datasets. M.J.R. and R.G. processed cryo-EM data. R.G. built atomic models. R.G. and K.H. reconstructed the tomograms. M.J.R. developed algorithms to denoise and segment tomograms, performed sub-tomogram averaging, and carried out hierarchical clustering / sub-tomogram variability analysis. K.R.C. and T.C.B. performed computational simulation studies. R.G., M.J.R., T.C.B., and G.M.A. wrote the paper with input from all other authors.

---

**Figure S1.**
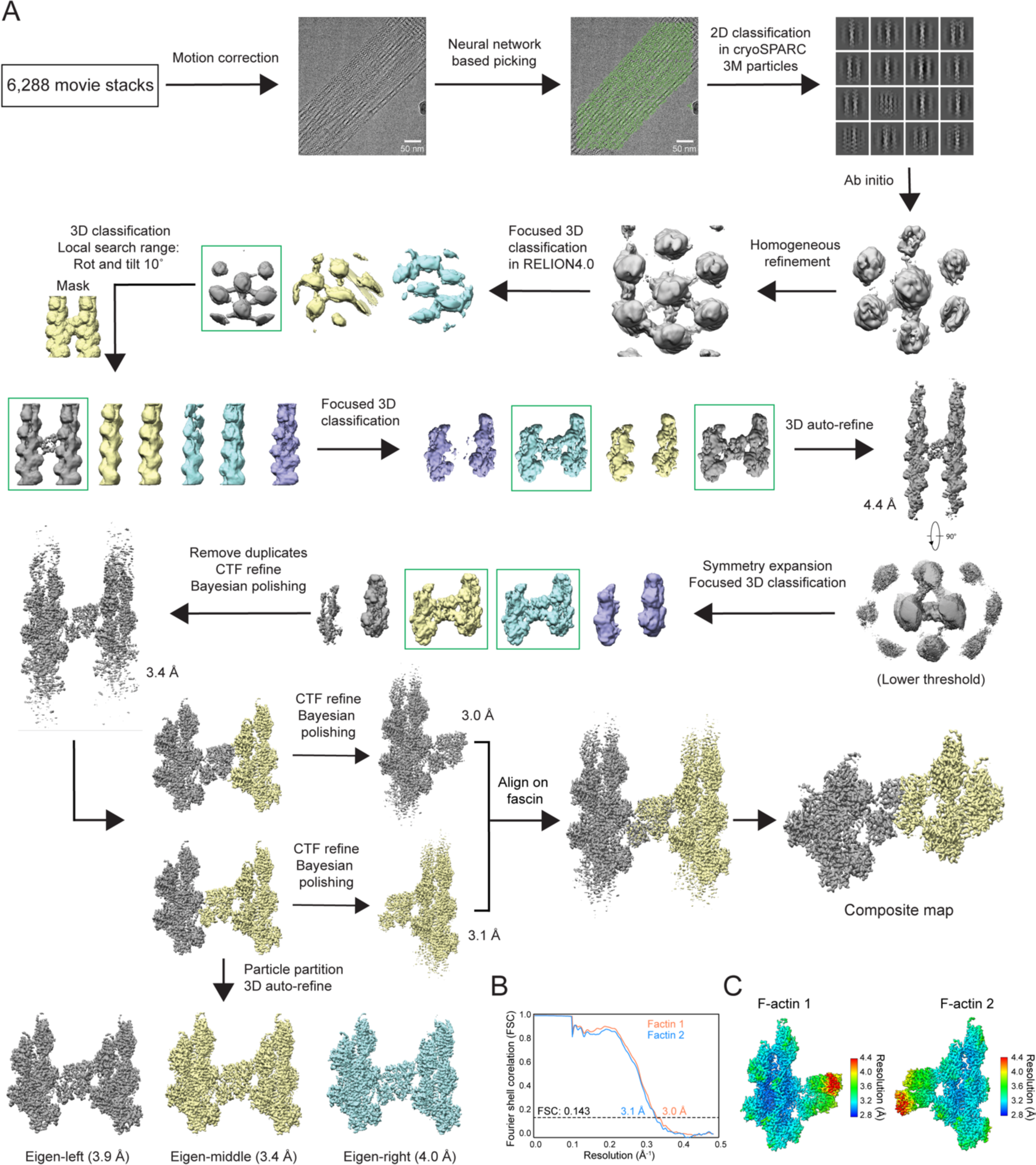
Cryo-EM data processing workflow. (A) Scheme of high resolution fascin crossbridge reconstruction data processing, including multibody refinement. (B) Fourier shell correlation (FSC) curves. (C) Local resolution assessment.

**Figure S2.**
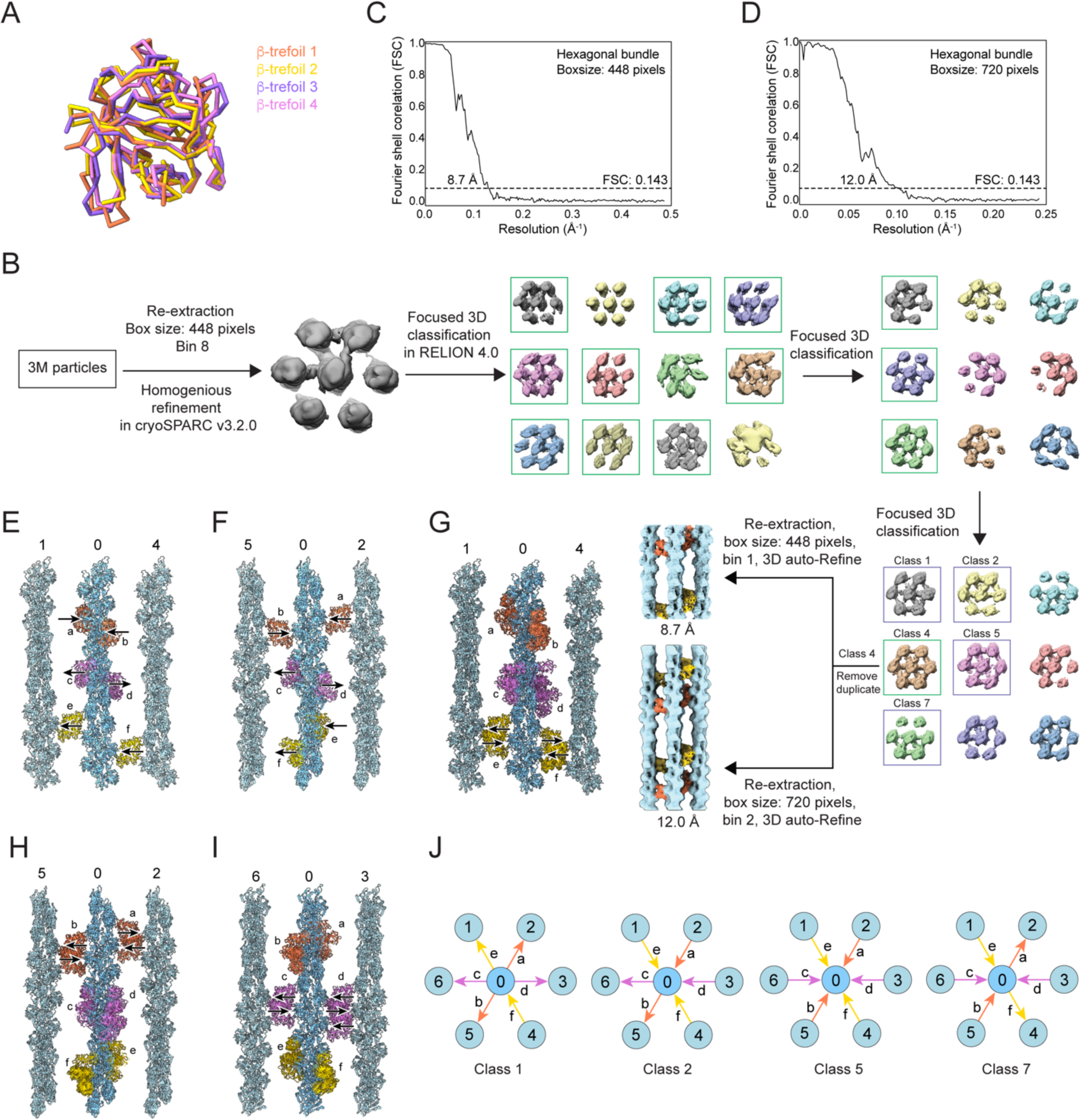
Analyses of fascin conformation and bundle element architecture. (A) Superposition of fascin’s four individual β-trefoil domains extracted from the consensus atomic model (Figure 1F). (B) Cryo-EM data processing workflow for reconstructing a hexagonal bundle element. (C-D) FSC curves for the bundle element reconstructed with two different box sizes. (E-F) Side views of actin planes 1-0-4 (E) and 5-0-2 (F) from the 8.7 Å reconstruction docking model. (G-I) Side view of each actin plane of models from 5 bundle element classes aligned on the central filament. (J) Distributions of poses of the 6 central-filament associated fascins from the four additional bundle element classes.

**Figure S3.**
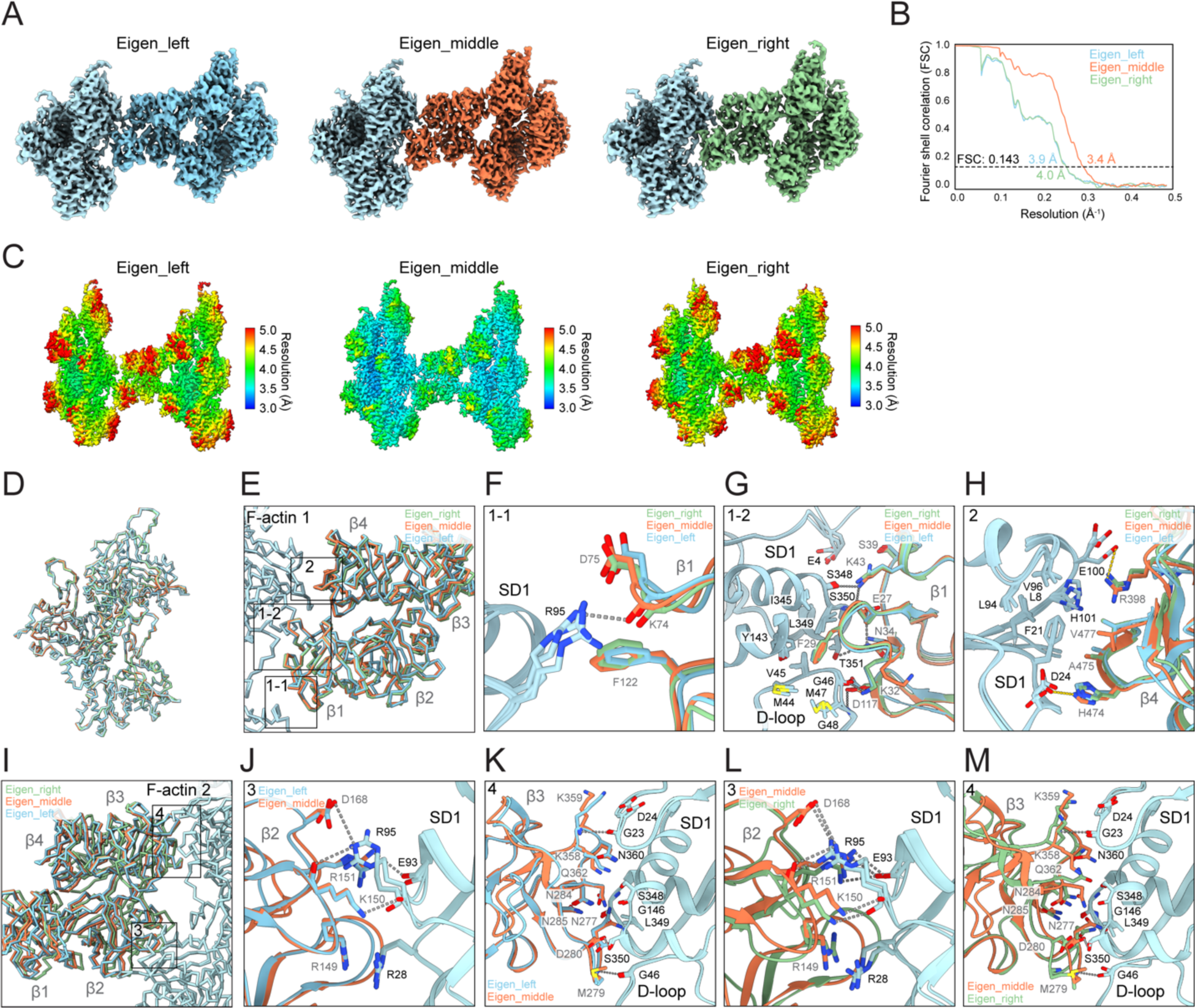
Analyses of multi-body derived reconstructions. (A) Cryo-EM density maps of eigen_left, eigen_middle and eigen_right reconstructions. (B-C) FSC curves (B) and local resolution assessment (C) of the three reconstructions. (D) Superposition of all six F-actin models from the three reconstructions. All F-actin 1 models are colored light blue, while F-actin 2 models are colored as in Figure 5B. (E) Comparison of the fascin-F-actin 1 interface across the three snapshots, superimposed on F-actin 1. (F-H) Detail views of contacts indicated in E. (I) Comparison of the fascin-F-actin 2 interface across the three snapshots, superimposed on F-actin 2. (J-M) Detail views of contacts indicated in I.

**Figure S4.**
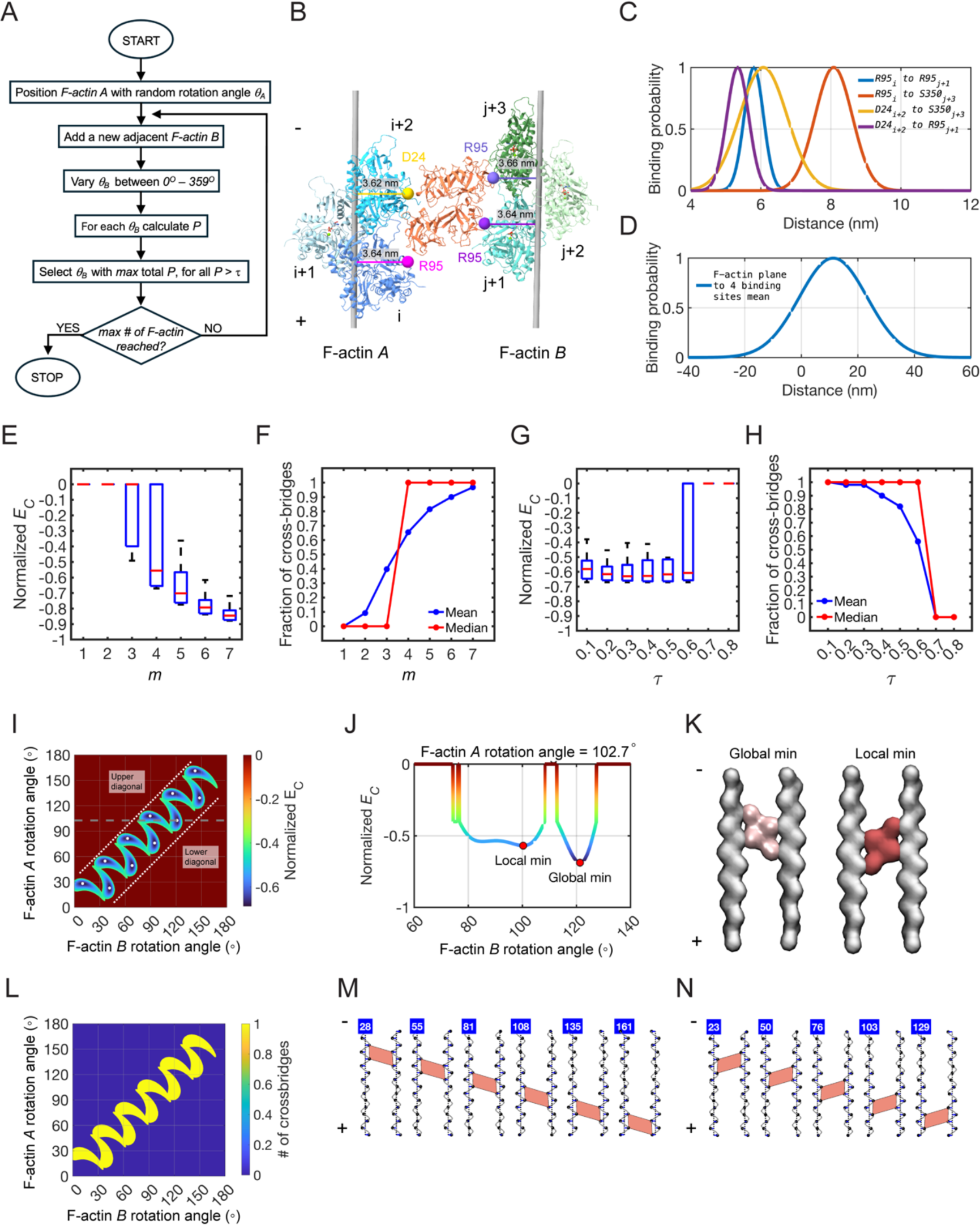
Computational model parametrization and analyses. (A) Flowchart of the computational model. Filaments are added iteratively, and the crosslinking probabilities for all angular rotations, *θB*, of the new filament *B* are evaluated. The angular rotation corresponding to the maximum probability of crosslinking is selected, as long as this probability is greater than the minimum cross-linking threshold, *τ*. (B) Atomic model of fascin crosslinked F-actin in ribbon representation. Filament *A* is depicted in shades of blue, while filament *B* is depicted in shades of green. Interfacial residues used as fiducials to calculate characteristic distances are indicated with spheres of varying colors. Grey cylinders represent F-actin axes. (C) Fascin binding probabilities as a function of distances between fiducial residues. (D) Fascin binding probability as a function of the separation between the geometric center of the four fiducial residues and the plane which spans both filament axes. (E) Distribution of normalized crossbridge energies for different values of the standard deviation multiplier, *m*, between 1 and 7. N = 2,800 independent runs of filament pairs. Filament *A* was initialized with random absolute rotation angles, while *τ* was varied systematically between 0.1 and 0.8 in increments of 0.1. Red line indicates median; lower and upper bounds of box are 25^th^ and 75^th^ percentile, respectively. (F) Reanalysis of simulation results from E, plotting the fraction of crossbridges. (G) Boxplot of the distribution of normalized crossbridge energies for different values of *τ* using *m* = 4. N = 400 independent runs of filament pairs. F-actin *A* was initialized with random absolute rotation angles. Red line indicates median; lower and upper bounds of box are 25^th^ and 75^th^ percentile, respectively. (H) Reanalysis of simulation results from G, plotting the fraction of crossbridges. (I) Heatmap of the normalized cross-bridge energies for a filament pair as a function of absolute rotation angles. White dots show energy minima. (J) Plot of normalized crossbridge energy for F-actin *A* rotation angle = 102.7° and varying the rotation angle of F-actin *B* between 60° and 140 °, with increments of 0.1°, corresponding to gray dashed line in I. (K) Surface representation of a fascin-crosslinked filament pair for the global and local minimum indicated in J. F-actins are represented in grey. For the global minimum, fascin is in pink and corresponds to *P* = 0.69. For the local minimum, fascin is in red, corresponding to *P* = 0.57. (L) Heatmap of the number of crossbridges for filament pairs with different rotational shifts. Each filament has 14 protomers. (M) Representative model renderings of the minima in the upper diagonal in I, showing fascin crossbridges in the “down” pose. (N) Representative model renderings of the minima in the lower diagonal in panel I, showing fascin crossbridges in the “up” pose. Blue boxes indicate filament *A* rotation angles.

**Figure S5.**
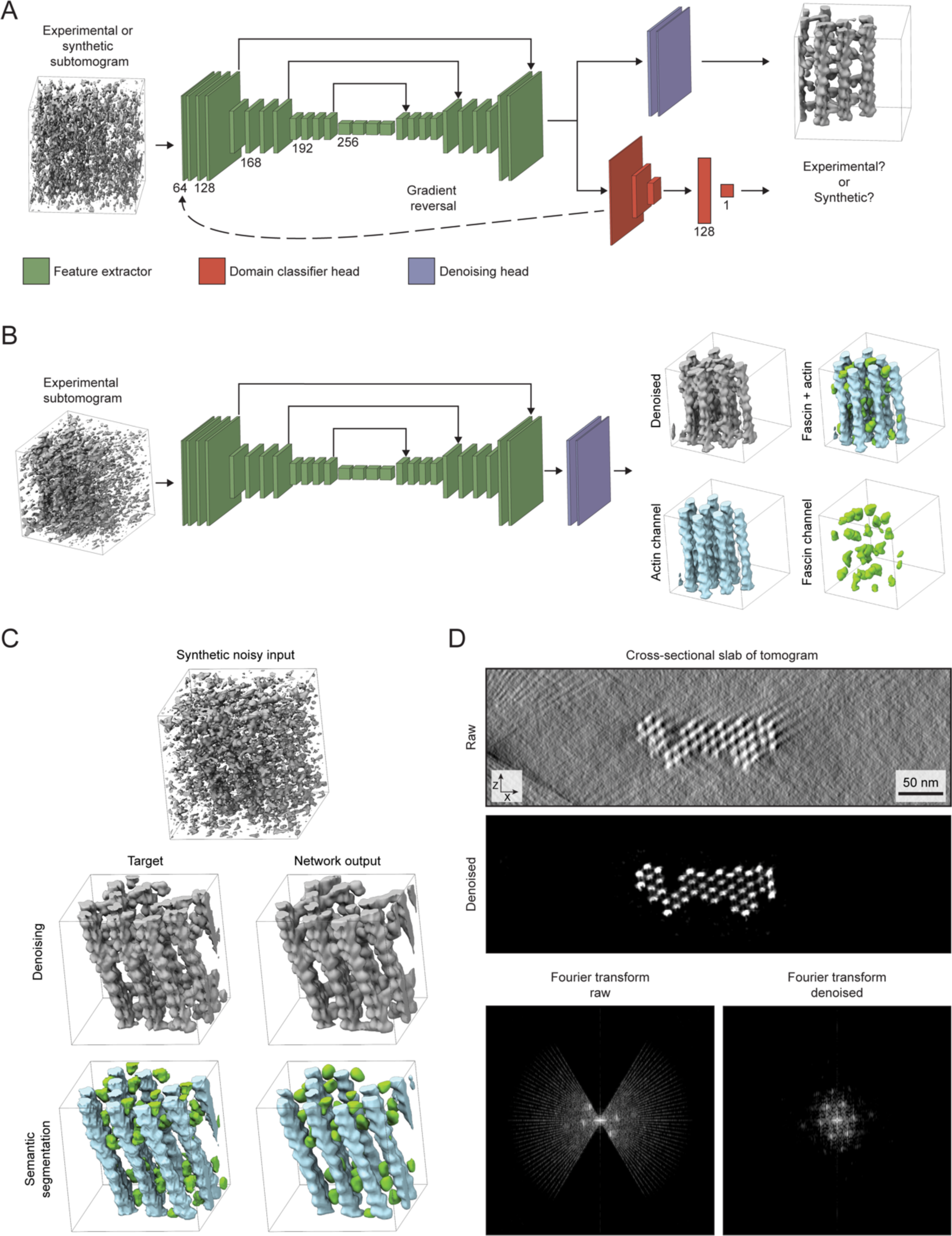
Neural network architecture and performance. (A) Neural network architecture used for pretraining on synthetic data and training on both synthetic and experimental subtomograms. (B) Neural network architecture and performance used for inference on experimental subtomogram. (C) Representative denoising and semantic segmentation performance on synthetic, noisy subtomogram. (D) Neural network denoising performance on cross-sectional slab of experimental tomogram.

**Figure S6.**
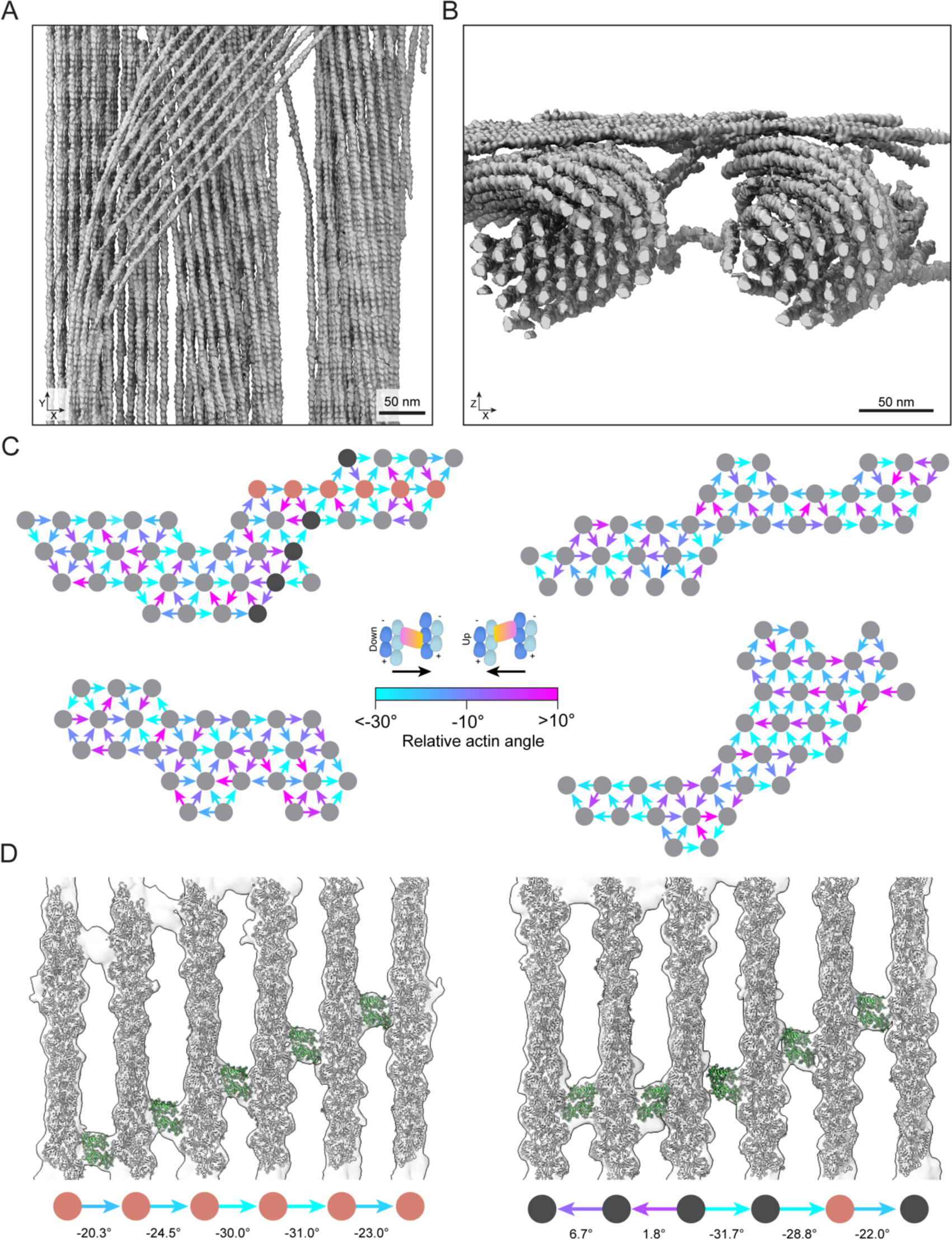
Additional structural analysis of fascin crosslinked bundles. (A) Top view of semantically segmented F-actin highlighting interconnected bundles. (B) End-on view of bundles in A, highlighting supertwist along their respective longitudinal axes. (C) End-on view schematics of filament rotational phase shifts and fascin poses of four additional bundles, analyzed as in Figure 6B. Dark grey and gold filaments correspond to actin planes displayed in D. (D) Side views of rigid-body docking models of two additional actin planes. Filament rotational phase offsets and fascin poses are indicated.

**Figure S7.**
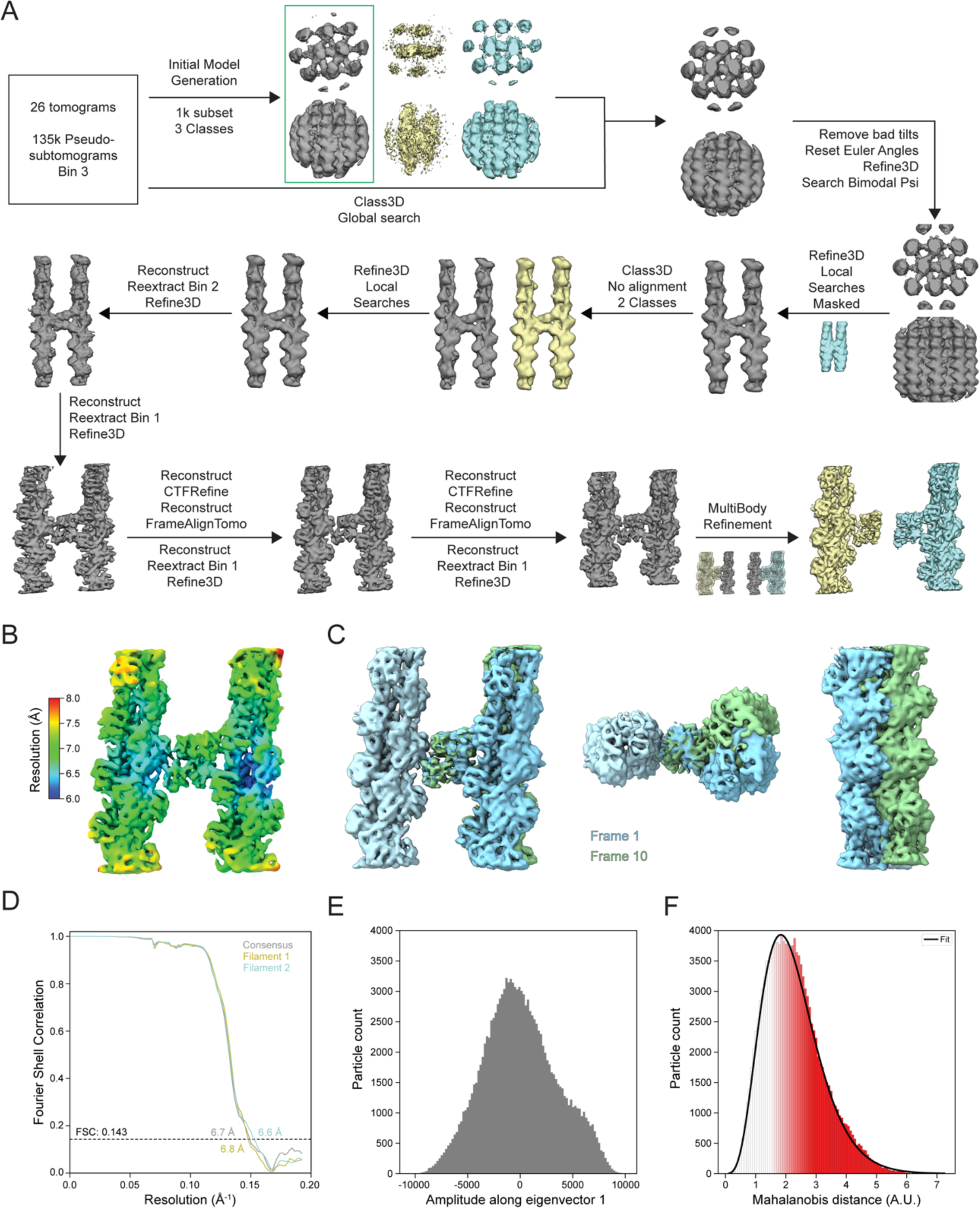
Subtomogram averaging workflow and analyses. (A) Subtomogram averaging data processing workflow. (B) Local resolution map of subtomogram average. (C) Extreme frames (1 representing 5^th^ percentile, and 10 representing 95^th^ percentile) of interpolation along the first principal component of multibody refinement. Similar inter-filament rotation is present as observed in single particle analysis. (D) FSC curves of the consensus (gray) and multi-body refinement reconstructions (yellow, blue). (E) Distribution of amplitudes along eigenvector 1 of all subtomograms. (F) Distribution of Mahalanobis distances of all subtomograms (n = 129,948). Fit represents gamma distribution (R^2^ = 0.9951).

**Table S1.** Cryo-EM data collection, refinement, and validation statistics.

| Data collection |  |  |  |  |  |  |  |  |
| --- | --- | --- | --- | --- | --- | --- | --- | --- |
| Microscope | Titan Krios |  |  |  |  |  |  |  |
| Voltage (kV) | 300 |  |  |  |  |  |  |  |
| Detector | K2 Summit |  |  |  |  |  |  |  |
| Magnification | 29,000 |  |  |  |  |  |  |  |
| Electron exposure (e<sup>-</sup> / Å<sup>2</sup>) | 61.26 |  |  |  |  |  |  |  |
| Exposure rate (e<sup>-</sup> / pixel / s) | 6.5 |  |  |  |  |  |  |  |
| Calibrated pixel size (Å) | 1.03 |  |  |  |  |  |  |  |
| Defocus range (µm) | −0.8 to −2.2 |  |  |  |  |  |  |  |
| Symmetry imposed | C1 |  |  |  |  |  |  |  |
| Data processing | Fascin crosslinked F-actin |  |  |  |  |  | Bundle element |  |
|  | Multibody: F-actin 1 | Multibody: F-actin 2 | Composite map | Eigen_left | Eigen_middle | Eigen_right | Box: 460 Å | Box: 740 Å |
| Initial particle images (no.) | 3,056,360 |  |  |  |  |  | 3,056,360 | 3,056,360 |
| Final particle images (no.) | 113,800 |  |  | 17,207 | 79,824 | 16,769 | 8,477 | 8,053 |
| Map resolution (Å) | 3.0 | 3.1 |  | 3.9 | 3.4 | 4.0 | 8.7 | 12.0 |
| FSC threshold | 0.143 |  |  |  |  |  |  |  |
| Refinement |  |  |  |  |  |  |  |  |
| Initial model (PDB ID) | 7R8V, 3LLP | 7R8V, 3LLP | 7R8V, 3LLP | 7R8V, 3LLP | 7R8V, 3LLP | 7R8V, 3LLP |  |  |
| Model resolution (Å) | 3.1 | 3.0 | 3.1 | 3.6 | 3.5 | 3.6 |  |  |
| FSC threshold | 0.5 | 0.5 | 0.5 | 0.5 | 0.5 | 0.5 |  |  |
| Map sharpening B factor (Å<sup>2</sup>) | -55.11 | -48.84 |  | -50.28 | -62.33 | -48.48 | -108.59 | -512.65 |
| Model composition | 3 actin protomers, 1 fascin | 3 actin protomers, 1 fascin | 6 actin protomers, 1 fascin | 6 actin protomers, 1 fascin | 6 actin protomers, 1 fascin | 6 actin protomers, 1 fascin |  |  |
| Non-hydrogen atoms | 12,625 | 12,625 | 21,460 | 21,460 | 21,460 | 21,460 |  |  |
| Protein residues | 1,604 | 1,604 | 2,723 | 2,723 | 2,723 | 2,723 |  |  |
| Ligands | 3 Mg.ADP | 3 Mg.ADP | 6 Mg.ADP | 6 Mg.ADP | 6 Mg.ADP | 6 Mg.ADP |  |  |
| B factors (Å<sup>2</sup>) |  |  |  |  |  |  |  |  |
| Protein | 23.94 | 27.87 | 43.13 | 99.43 | 78.72 | 112.12 |  |  |
| Ligand | 8.28 | 18.9 | 39.23 | 90.45 | 65.04 | 96.87 |  |  |
| R.M.S. deviations |  |  |  |  |  |  |  |  |
| Bond lengths (Å) | 0.003 | 0.005 | 0.004 | 0.004 | 0.002 | 0.004 |  |  |
| Bond angles (°) | 0.583 | 0.663 | 0.612 | 0.667 | 0.566 | 0.614 |  |  |
| Validation |  |  |  |  |  |  |  |  |
| MolProbity score | 1.55 | 1.73 | 1.55 | 1.94 | 1.63 | 2.07 |  |  |
| Clash score | 7.37 | 8.65 | 8.07 | 16.31 | 8.24 | 18.66 |  |  |
| Poor rotamers (%) | 0 | 0 | 0 | 0.04 | 0.07 | 0.09 |  |  |
| Ramachandran plot |  |  |  |  |  |  |  |  |
| Favored (%) | 97.23 | 96.16 | 97.44 | 96.58 | 96.92 | 95.62 |  |  |
| Allowed (%) | 2.77 | 3.84 | 2.56 | 3.42 | 3.08 | 4.38 |  |  |
| Disallowed (%) | 0.00 | 0.00 | 0.00 | 0.00 | 0.00 | 0.00 |  |  |

**Table S2.** Cryo-ET data collection and subtomogram averaging.

| Data collection |  |  |  |
| --- | --- | --- | --- |
| Microscope | Titan Krios |  |  |
| Voltage (kV) | 300 |  |  |
| Detector | K3 |  |  |
| Magnification | 26,000 |  |  |
| Electron exposure (e<sup>-</sup> / Å<sup>2</sup>) | 109.17 |  |  |
| Exposure rate (e<sup>-</sup> / pixel / s) | 30 |  |  |
| Calibrated pixel size (Å) | 2.6 |  |  |
| Defocus (μm) | -4.0 |  |  |
| Symmetry Imposed | C1 |  |  |
| Data Processing | Multibody: F-actin 1 | Multibody: F-actin 2 | Composite map |
| Initial particle images (no.) | 134,733 |  |  |
| Final particle images (no.) | 129,948 |  |  |
| Map resolution (Å) | 6.8 | 6.6 | 6.7 |
| FSC threshold | 0.143 | 0.143 | 0.143 |
| Map sharpening B factor (Å<sup>2</sup>) | -301.274 | -280.458 | -233.239 |

**Table S3.**
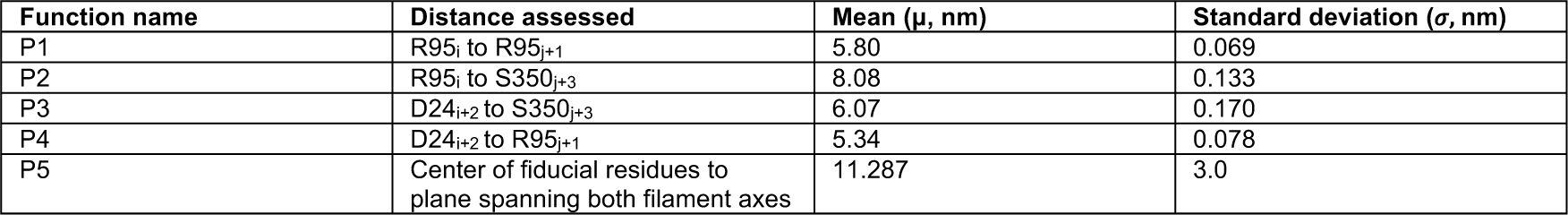
Computational modeling probability function parameters.

## Supplementary Video Captions

**Video S1: Morphs between prebound, inhibitor bound, and F-actin bound fascin conformations**

Structures are superimposed on fascin β-trefoil 2.

**Video S2: Overview of fascin crosslinked F-actin bundle element architecture**

**Video S3: Morph between eigen_left, eigen_middle, and eigen_right conformational snapshots**

Structures are superimposed on F-actin 1.

**Video S4: Overview of a tomogram showcasing denoising and semantic segmentation**

## Methods

### Protein expression and purification

The coding sequence of human fascin1 (GeneBank, NM_003088.4) was obtained from Addgene (#31207) and subsequently inserted into a pGEX-6p-1 vector (Cytiva, 28954648) using Gibson Assembly (New England Biolabs, E2611L). The resulting construct was expressed in *Escherichia coli* BL21(DE3) cells (New England Biolabs, C2527H) at 18 °C for 16 hours after induction with 0.2 mM isopropyl-β-D-thiogalactoside (Goldbio, I2481C100). The cells were resuspended and disrupted in lysis buffer (25 mM Tris-HCl pH 8.0, 150 mM NaCl). After centrifugation at 20,000 rpm for 30 minutes in a JA-25.50 rotor (Beckman Coulter), GST-tagged fascin in the supernatant was enriched using GST affinity beads (Glutathione Sepharose 4 Fast Flow, Cytiva). The GST tag was removed by incubating the beads with 3C protease^1^ at 4 °C overnight. The cleaved target protein was purified on a HiTrap Q HP ion exchange column (Cytiva), and further polished on a Superdex 200 10/300 Increase column (Cytiva) equilibrated with 25 mM Tris-HCl pH 8.0, 150 mM NaCl and 3 mM dithiothreitol (DTT).

Chicken skeletal muscle actin was purified as previously described^2^ and stored at 4 °C in G-Ca buffer: 2 mM Tris-HCl pH 8.0, 0.5 mM DTT, 0.1 mM CaCl_2_, 0.2 mM ATP, 0.01% NaN_3_. F-actin was polymerized at room temperature for 1 hour by mixing 5 μM monomeric actin in G-Mg buffer (2 mM Tris-HCl pH 8.0, 0.5 mM DTT, 0.2 mM ATP, 0.1 mM MgCl_2_) with KMEI buffer (50 mM KCl, 1 mM MgCl_2_, 1 mM ethylene glycol-bis(β-aminoethyl ether)-N,N,N′,N′-tetraacetic acid, 10 mM imidazole pH 7.0) supplemented with 0.01% Nonidet P40 (NP40) substitute (Roche). The polymerized F-actin was stored at 4 °C overnight before grid preparation.

### Cryo-EM and cryo-ET sample preparation

To prepare fascin crosslinked F-actin bundle for cryo-EM analysis, 0.6 μM F-actin was incubated with fascin at a molar ratio of 1:2 at room temperature for 30 minutes. For cryo-ET analysis, F-actin bundles were prepared by incubating 0.3 μM each of F-actin and fascin at room temperature for 72 hours. 4 μl of the specimen was applied to a freshly plasma cleaned (Gatan Solarus, H_2_ / O_2_ mixture) C-flat 1.2/1.3 holey carbon Au 300 mesh grid (Electron Microscopy Sciences) in a Leica EM GP plunge freezer operating at 25 °C and 100% humidity. After 1 minute of incubation, the grid was blotted from the back with a Whatman no. 5 filter paper for 4 s and plunge-frozen in liquid ethane cooled by liquid nitrogen.

### Cryo-EM and cryo-ET data acquisition

Cryo-EM data were collected on a Titan Krios microscope (FEI), operating at 300 kV and equipped with a Gatan K2-summit detector in super-resolution mode using SerialEM^3^. Frame sequences (movies) were recorded at a nominal magnification of 29,000X, corresponding to a calibrated pixel size of 1.03 Å at the specimen level (super-resolution pixel size of 0.515 Å / pixel). Each exposure was fractionated across 40 frames with a total electron dose of 61 e^-^ / Å^2^ (1.53 e^-^ / Å^2^ / frame) and a total exposure time of 10 s. The defocus values ranged from -0.8 to -2.2 μm. To alleviate the effects of potential preferential orientation of fascin within F-actin bundles, data were collected with the stage tilted at three different angles: 30°, 15°, and 0°. At 30° and 15°, exposures were acquired by targeting a single hole per stage translation. At 0°, exposures were acquired using the beam tilt / image shift strategy, targeting 9 holes per stage translation. A total of 6,288 micrographs were collected, with 1,258, 1,439, and 3,591 micrographs obtained at 30°, 15°, and 0°, respectively.

Cryo-ET data were collected on a spherical-aberration (C_s_) corrected Titan Krios microscope operating at 300 kV, equipped with a Gatan K3 detector and BioQuantum energy filter (slit width 20 eV). Tilt series were recorded from -60° to 60° with a tilt increment of 3°. The image stack at each tilt angle was acquired at a nominal magnification of 26,000x corresponding to a calibrated pixel size of 2.6 Å at the specimen level (super-resolution pixel size of 1.3 Å / pixel). Each stack was fractionated into 12 frames with a total electron dose of 2.66 e^-^ / Å^2^ (0.22 e^-^ / Å^2^ / frame) and a total exposure time of 0.6 s. All 27 tilt series were collected at a nominal defocus of -4 μm.

### Cryo-EM image processing

Frame sequences were motion corrected, dose weighted and summed with 2 x 2 binning (to a 1.03 Å pixel size) using MotionCor2^4^. Contrast Transfer Function (CTF) estimation was performed with CTFFIND4^5^ using non-dose weighted sums. Particle picking was performed with a previously reported neural-network–based approach we developed for handling F-actin bundles^6^. Briefly, PDB 7R8V was expanded along its helical axis to produce a filament with 49 protomers and converted to a volume using the molmap command in Chimera^10^. Synthetic data was procedurally generated by spawning bundle pairs from two of copies of this volume featuring a horizontal displacement randomly sampled from a Gaussian distribution centered at 113.8 Å with a standard deviation of 15.9 Å (estimated by manually measuring the inter-filament distances of 36 bundles in our dataset) and a vertical displacement randomly sampled from a uniform distribution between -180 Å and +180 Å. A random angular skew sampled from a Gaussian distribution with a mean of 0° and a standard deviation of 1.5° was also applied to the second filament.

To generate synthetic images, 0,1,2, or 3 of these bundle pairs were loaded while applying a translation in X, Y, and Z each randomly sampled from a uniform distribution between -250 Å to +250 Å. This composite volume was then projected along the Z dimension to generate a noiseless projection. The first (rot) and third (psi) Euler angles were randomly sampled from a uniform circle in degree increments and the second (tilt) Euler angle was randomly sampled from a Gaussian distribution with a mean of 0° and a standard deviation of 7.5°. These projections were corrupted by a theoretical contrast transfer function and pink noise using EMAN2 functions^7^. To generate training data for semantic segmentation, the noiseless projections were lowpass-filtered to 40 Å, a binarization threshold of 0.9 was applied, and morphological closing was performed by first dilating by 66 Å then eroding by 33 Å. A denoising autoencoder neural network featuring the same architecture as previously described^6^ was trained using 150,000 noisy and noiseless projection pairs and a 90 : 10 training : validation split with a learning rate of 0.00005 until convergence after 14 epochs with a validation cross-correlation coefficient loss of 0.9296. This pre-trained network was then trained for semantic segmentation using 50,000 projection pairs and a 90 : 10 training : validation split with a learning rate of 0.00005 until convergence after 13 epochs with a validation categorical cross-entropy of 0.787. The trained neural network for semantic segmentation was then used for inference on the experimental images to identify pixels containing F-actin bundles. Tiles of 192 pixels spaced in 48 pixel increments from micrographs binned by 4 were passed through the neural network for semantic segmentation before being stitched back together via a maximum intensity operation. From these semantic maps, particle picks were generated by removing overlap within a 60 Å distance.

The coordinates of 3,056,360 picked particles were imported into RELION4.0^8^ and extracted with a box size of 448 pixels, then binned by 2. The extracted particles were then imported into cryoSPARC v4.2^9^ for reference-free 2D classification. 2,231,774 particles were selected from classes featuring parallel actin filaments. A quarter of them (550,000) were used for *ab initio* 3D reconstruction to generate an initial reference. This reconstruction featured seven filaments arranged in a hexagonal lattice with several apparent inter-filament densities corresponding to fascin crossbridges (Figure S1A). The full selected set of 2,231,774 particles were then subjected to masked homogeneous refinement using a cylinder-shaped mask covering all 7 filaments. The aligned particles were subsequently re-imported into RELION for 3D classification without image alignment. Particles from the class displaying the clearest hexagonal arrangement were selected, re-centered on the fascin cross-bridge with the strongest density, re-extracted, and reconstructed. An “H” shaped mask covering the re-centered fascin cross-bridge and its two associated filaments was created. The re-centered particles were then subjected to masked 3D classification with a global search range for rot angles and a 10° local search range for psi and tilt angles. Particles with clear cross-bridge density were selected and subjected to another round of focused 3D classification with a mask covering the fascin crossbridge and its two interacting actin subunits on each filament. Classes with improved fascin density were pooled and refined using the “H” shaped mask, yielding a density map displaying strong fascin density within the mask and four neighboring weak fascin densities.

Particles were then subjected to symmetry expansion with re-centering on each of the additional fascin densities, followed by focused 3D classification. Particles corresponding to classes with strong crossbridge density were then re-extracted without binning at a pixel size of 1.03 Å. After duplicate removal, CTF-refinement, Bayesian polishing, 3D refinement, and postprocessing, we obtained a consensus 3.4 Å resolution density map featuring streaking artifacts in fascin (Figure S1). To further improve the map quality, two rounds of multibody refinement were performed, alternately masking fascin and one filament as one body and the other filament as the other body (i.e., F-actin 1 + fascin as body 1 and F-actin 2 as body 2; then F-actin 2 + fascin as body 1 and F-actin 1 as body 2). This greatly improved the density of each F-actin-fascin interface when it was within the body 1 mask. The final 3D refinements yielded postprocessed density maps with resolutions of 3.1 Å and 3.0 Å, respectively.

To generate a composite map containing both well-resolved fascin–F-actin interfaces, the two multi-body maps were aligned on fascin. The poorly resolved pair of β-trefoil domains from each map was removed with the “split map” command, then the remaining well-resolved density from the two maps was stitched together using the “vop maximum” command, both in UCSF Chimera^10^.

To resolve a fascin crosslinked F-actin hexagonal bundle element, the original 3,056,360 picked particles were extracted in RELION in a box size of 448 pixels and binned by 8. Particles were then imported into Cryosparc and aligned by homogeneous refinement with a cylinder-shaped mask and a hexagonal bundle reference generated from the abovementioned *ab initio* 3D reconstruction. Aligned particles were subjected to several rounds of focused 3D classification in RELION to enrich particles with strong densities for all possible fascin molecules and F-actin filaments within a hexagonal bundle element. The 8,477 particles contributing to the class featuring the best density were re-extracted with a box size of 448 pixels and no binning, as well as a box size of 720 pixels binned by 2. Both particle stacks were reconstructed and refined, generating postprocessed density maps at 8.7 Å and 12.0 Å resolution, respectively. Four additional classes were refined which featured high-quality density for the central filament and discernable density for its six bound fascins, but varied quality for the peripheral filaments and crossbridges.

Local resolution estimation was performed using the procedure implemented in RELION.

### Tilt series reconstruction

Motion correction of individual frames and CTF estimation of combined tilt stacks were performed in WARP^11^. Data were then exported for subsequent alignment and reconstruction using the IMOD software package^12^. Tilt series were binned by 3 to a final pixel size of 7.8 Å, aligned with the patch-tracking approach, then reconstructed using weighted back-projection with a SIRT-like filter equivalent to 12 iterations.

### Synthetic tomography data generation

Synthetic datasets approximating subtomograms were generated to train neural networks for denoising and semantic segmentation. The training datasets for the neural networks consisted of noisy source volumes simulating empirical data, along with paired target volumes of *in silico* generated ground truth fascin–F-actin bundles. To produce these volumes, the seven-filament bundle element reconstruction was laterally expanded to produce a larger hypothetical lattice of 19 filaments bound by 84 fascins. Atomic models of F-actin (PDB 7R8V) and fascin crossbridges were fit into this extended volume, then converted into volume data using the molmap command in Chimera. These docked components were then used to generate plausible regions of bundles with varying architectures. For each simulated region, a random subset of the F-actin volumes was loaded. Fascins were then loaded only if they were in a position to bind these filaments; to simulate sub-stoichiometric binding, a random subset of these fascins were rejected. All of the F-actin volumes were then combined into a single volume, all of the fascin volumes were combined into a separate volume, and both volumes were saved. A library of 200 volume pairs was procedurally generated using this approach.

The library was then used to generate a dataset of 128 voxel training volumes at a sampling of 7.8 Å / voxel. Paired fascin and F-actin volumes were rotated about the rot and psi angles by random, uniformly sampled values between 0° and 359°, while the tilt angle was randomly sampled from a Gaussian probability distribution centered at 90° with a standard deviation of 20°. The density was then translated in the box along each dimension by a random uniform translation within the range of ±195 Å.

20,000 synthetic noisy volumes were generated by first projecting the repositioned volume at fixed angles corresponding to the experimental tilt series collection scheme (-60° to +60° with a 3° increment). These projections were corrupted by the CTF, then reconstructed back into a three-dimensional volume using reconstructor class functions as implemented in the EMAN2 python package^7^. Empirical noise was then extracted from each of our experimental tomograms by computing the average Fourier transform of 50 randomly sampled 128 voxel boxes. For each synthetic volume, one of these empirical noise boxes was randomly selected, multiplied element-wise by a white noise box of the same dimensions, normalized, and scaled by a random scale factor that modulated the signal-to-noise ratio of each synthetic particle. The synthetic volume was then summed with its noise volume in Fourier space. To account for interpolation artifacts from CTF application or noise addition in Fourier space, the synthetic volumes were then cropped to 64 voxels in real space. Each noisy particle had a corresponding noiseless ground truth particle, as well as binarized semantic maps calculated from the paired F-actin and fascin volumes.

### Neural network training for tomogram denoising and semantic segmentation

Pre-training of a denoising autoencoder consisting of 3D convolutional layers in a U-net architecture (Figure S5) was performed using a single NVIDIA A100 GPU with 80 GB of VRAM, using a learning rate of 0.0001. Training was run on the 20,000 pairs of noisy and ground truth volumes with a 90 : 10 training : validation split, until the network converged with a cross-correlation coefficient validation loss of 0.8659 after 20 epochs. Initial inference testing on experimental tomograms resulted in distorted denoised volumes. We reasoned this was because the synthetic data did not model corruption of the experimental data with sufficient fidelity. We therefore continued to train the network using a domain adversarial neural network (DANN) approach^13^. An additional network head was added for domain classification by forking the network output after the feature extraction layers (Figure S5A). The domain classification head consists of a gradient reversal layer and additional 3D convolutional layers, which are followed by flattening to a dense layer, then a binary classification layer with a sigmoid activation function. 100 synthetic volumes and 100 volumes extracted from the experimental tomograms were used as the training set for the domain classifier. Adversarial training was performed by alternatively passing these data through the domain classifier head, followed by re-training the feature extractors with the denoising head using only the 100 synthetic volumes. This adversarial training was run for ten iterations; however, after multiple iterations, too many high-resolution details were lost, so the first iteration of adversarial training was used for denoising.

After using DANN to complete training of the denoising autoencoder, a semantic segmentation network was trained using 10,000 volume sets and the autoencoder’s pre-trained weights. The final layer was adapted to produce multi-channel outputs, featuring a softmax activation layer and random initial weights. After training for 10 iterations with a learning rate of 0.00001, the categorical cross-entropy loss of the semantic segmentation network was 0.0556.

### Tomogram denoising and semantic segmentation

The trained neural networks were then used to denoise and semantically segment empirical tomograms, once again using single A100 GPUs. 64-voxel tiles sampled every 32 voxels in X and Y and every 4 voxels in Z were extracted, normalized, and passed as inputs to the neural network. The network outputs were masked with a 48-voxel cubic mask to minimize edge artifacts and stitched together via maximum intensity projection. Each input tomogram produced three outputs: a denoised tomogram, a semantic map of actin filaments, and a semantic map of fascin molecules.

### Measurement of F-actin rotational phase offsets in denoised tomograms

The denoised tomograms were manually inspected to identify bundles with high-quality density, straight filaments, and uniform fascin crossbands spanning multiple crossovers. Subsequently, a 17-subunit F-actin model was generated from PDB 7R8V and fit into filament densities in the bundle using Chimera while maintaining identical axial register of the filaments across the bundle. Synthetic density maps were then generated for each fitted F-actin atomic model using the “molmap” command in Chimera with a 6 Å resolution cutoff. The rotational phase offset was then measured by fitting the synthetic density map of the rotating filament to the reference filament using Chimera.

### Subtomogram averaging

Subtomogram particle picking required minimal post-processing of the fascin semantic maps. A threshold of 0.9 was applied to each semantic map, and the centroids of objects larger than 50 voxels were designated as potential fascin picks. The center of the hole in the carbon film for each tomogram was manually picked, and potential fascin picks within a 740-voxel radius of the hole center were retained as picks. Subtomogram averaging was performed using RELION4.0. From 26 tomograms, 135,000 192-voxel pseudo-subtomograms were extracted at bin 3 (voxel size 7.8 Å) and cropped to a 64-voxel box. A small subset of 1,000 pseudo-subtomograms was used for initial model generation with three classes. Two of the three classes contained 82% of the particles, and the most populous class was selected. An initial global alignment of particles was performed using Class3D with one class. 5,000 particles with tilt angles outside of the range 60 to 120° were subsequently excluded. The rot angle was then removed from the metadata, and a bimodal psi prior was used for subsequent 3D auto-refinement. An H-shaped mask (85% z-length) was used for focused 3D auto-refinement and subsequent processing of the central filaments and fascin bridge. 3D classification using two classes was attempted to remove partially decorated fascin monomers or picks of single filaments, but both classes were essentially identical, suggesting a minimal false-positive rate in picking.

After another round of local 3D auto-refinement, pseudo-subtomograms were then sequentially re-extracted and subjected to local 3D auto-refinement at bin 2, then bin 1. This produced a reconstruction with a nominal resolution of 8.5 Å, yet the map appeared distorted. After two rounds of CTF refinement, FrameAlignTomo, and local 3D auto-refinement, a final undistorted 6.7 Å reconstruction was obtained. Multi-body refinement was then conducted in an identical manner as described above for the single particle analysis. Local resolution estimation was performed using RELION’s implementation.

### Bundle hierarchical clustering based on subtomogram variability analysis

Per-subtomogram deviations from the consensus reconstruction were measured using the Mahalanobis distance of each particle’s multi-body refinement amplitude along the eigenvectors corresponding to rotations. To calculate filament interface scores, fascin crossbridges were assigned to inter-filament interfaces using a custom script. The filament interface score (F.I.S) was computed as:

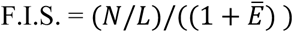

where *N* is the number of fascin crossbridges along the interface, *L* is the length of the interface, and *E*^)^ is the average energy (Mahalonbis distance) of the crossbridges along the interface. Graphs were constructed from 24 well-ordered bundle regions approximately 400 nm in length, where actin filaments correspond to nodes, and nodes are joined by an edge if the corresponding actin filaments were bridged by at least one fascin within the region. Edge weights were assigned as the filament interface scores.

Hierarchical clustering was performed using the ward linkage method, and the maximum distance was normalized across all graphs to enable direct comparisons. The metrics of transitivity, modularity, and minimum, average, and maximum cluster sizes were computed as a function of linkage distance for each graph.

### Atomic model building, refinement, and analysis

Previously determined structures of actin (PDB: 7R8V)^2^ and fascin (PDB: 3LLP)^14^ were rigid-body docked into each of the high resolution multi-body refinement derived density maps in ChimeraX v1.6.1^15^, then flexibly fitted with ISOLDE^16^. Atomic models were then refined using phenix.real_space_refine^17^ alternating with manual adjustments in Coot^18^. Refined models were then rigid-body fit into the composite map, split at the boundary between the well-resolved and poorly-resolved β-trefoil domains in the multi-body map from which the model was derived, merged, and real-spaced refined. Model validation was conducted with MolProbity^19^ as implemented in Phenix. The structural rearrangements of fascin in different states were analyzed with the DynDom online server^20^.

### Computational model

A computational model that iteratively optimizes the geometry of actin filaments and fascins based on multi-body refinement analysis was developed to assess their three-dimensional arrangement in hexagonal arrays. The model considers double-stranded helical F-actins as linear chains of 14-19 monomers, with each monomer rotated by -166.7 and translated by 2.75 nm, along the vertical axis. The axis-to-axis distance between F-actins in the hexagonal array bundle is 12.15 nm.

### Geometry initialization and algorithm implementation

The algorithm constructs an array of filaments iteratively until a maximum filament number is reached, and at each iteration it maximizes the probability, *P*, of forming fascin cross-bridges between the existing filaments and the newly incorporated filament (Figure S4A). Each filament is identified by its absolute rotation angle, corresponding to the angle that the first monomer forms with the origin of the simulation domain. The algorithm starts by positioning the first filament, *A*, at a random position in the bundle and with a random absolute rotation angle *θ_A_*. In the first iteration, it adds a second filament, *B*, and evaluates the probability of forming a cross-bridge, *P*, for each possible rotation angle of filament *B, θ_B_*, between 0-359.9° in increments of 0.1°. Last, it selects *θ_B_* that maximizes *P*, given that *P* > *τ*, the minimum threshold for fascin binding (Figure S4A). After the second filament is added, the following filaments are added within 12.15 nm of at least two existing F-actins. Actin protomers are allowed to form a maximum of one fascin bond, which remains static for the remainder of the simulation.

### Scoring functions

For each pair of filaments *A* and *B*, five distances are calculated to evaluate crossbridging probability between the two filaments and the position of the crossbridge. Considering all pairs of two protomers, *i* and *i+2* from filament *A* and *j+1* and *j+3* from filament *B*, the probability of an “up” pose fascin bond, *P*, is defined as the product of five probabilities. A fascin crossbridge is formed between protomers *i* and *i+2* from filament *A,* and protomers *j+1* and *j+3* from filament *B* for the largest *P* if that *P* is greater than *τ*. The probabilities of “down” pose bonds are simultaneously calculated by the algorithm by assigning the index *j* to filament *A* and the index *i* to filament *B*.

The five cross-bridge probabilities, *P_1,2,3,4,5_*, used to calculate *P* are based on four pairwise distances between fiducial residues corresponding to fascin-actin contact sites (Figure S4B) and one distance between the center of these sites and a plane defined by the axes of the filaments, which is included to prevent unrealistic fascin distortions. The four pairwise distances correspond to the separation between fascin binding sites on four protomers: R95_i_ and R95_j+1_; R95_i_ and S350_j+3_; D24_i+2_ and S350_j+3_; and D24_i+2_ and R95_j+1_ (Figure S4B).

Considering *n = 1, 2, …, 5* for the five different cross-bridging probabilities, the algorithm calculates the individual distance-dependent probabilities as:

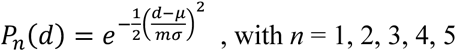

where *d* is the distance, while *μ* and *σ* are the mean and standard deviations of *d*, respectively. Their values are extracted from the multibody refinement analysis (Table S3). The parameter *m* is the multiplication factor for the standard deviation, *σ*, which effectively encodes fascin plasticity as it modulates the range of pairwise distances with high probability. The normalized cross-bridge energy is calculated as the negative inverse of *P*: 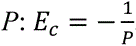, such that higher *P* corresponds to lower *E_c_*. Distributions of these probabilities versus d are presented in Figures S4C and S4D.

### Model parameterization

The two free model parameters, *m* and *τ*, were adjusted based on systematic analysis of their effects on the distribution of *P* and on the fraction of filament pairs which formed a fascin crossbridge. Filaments pairs with randomized angular shifts were used for this analysis. First, we tested how systematic variations of *m* affected the distribution of *P*. For each *m* between 1-7 (step size 1), and at each different value of *τ* from 0.1 - 0.8 (step size 0.1), 50 independent runs were performed, for a total of 2,800 runs. The normalized crossbridge energy distribution features a biphasic relation with *m*: it first increases as *m* increases from 1 to 4, then it decreases with *m* > 4 (Figure S4E). Next, we tested how systematically varying *m* affects the fraction of crossbridges formed across runs. We observed a low fraction of crossbridges for *m* < 4, with an average cross-bridge fraction < 0.5 and a median fraction of 0 (Figure S4F). Based on these data, a specific value of *m* was able to capture variability in the probabilities of crosslinking, with a high enough fraction of resulting fascin bonds: *m* = 4 (Figure S4E and S4F). The sensitivity of the model outputs was also assessed for *τ*. For these simulations, *m* was fixed at 4 as *τ* was varied between 0.1 and 0.8 (stepsize of 0.1), with 50 model runs with randomized angular shifts per *τ* step, for a total of 400 independent runs. For *τ* > 0.6, *E_c_* = 0, meaning that *τ* was too restrictive to allow crossbridges to form (Figure S4G). For *τ* ≤ 0.6, the distribution of *E_c_* broadened, providing a range (0.1 ≤ *τ* ≤ 0.6) that produced fascin crossbridging (Figure S4G). Analysis of the mean and median crossbridging fraction further revealed that, within that range, using *τ* < 0.4 was too permissive as it allowed constitutive bond formation (Figure S4H). Hence, the range of acceptable *τ* values was identified as 0.4 ≤ *τ* ≤ 0.6. Based on this range, *τ* = 0.4 was selected for further simulations of higher-order fascin crosslinked assemblies.

We first analyzed pairs of filaments composed of 14 protomers (spanning a single crossover) to evaluate the effect of relative angular shifts on the probability fascin cross-bridging, indicated as normalized energy, *E_c_*. Systematic variation of the two filaments’ rotation angles between 0-180° identify regions without crossbridging, corresponding to *E_c_* = 0, and regions of crossbridging with values of normalized *E_c_* < 0 (Figure S4I). Consistent with the periodicity of F-actin, we observe a regular pattern of teardrop-shaped energy wells around minima (indicated by white dots in Figure S4J). To explore the implications of this pattern, we first examined *E_c_* with the absolute rotation angle of filament *A* held constant at 102.7° while rotating filament *B* between 60-140°. This revealed one global minimum in *E_c_* and several local minima (Figure S4J). We find that in this case the global minimum corresponds to a fascin bridge in the “down” pose, while the most favorable local minimum corresponds to fascin bridge in the “up” pose at a nearly equivalent position (Figure S4K). This suggests the two teardrops which intersect a given filament *A* angle correspond to nearly isoenergetic “up” and “down” binding positions, whose relative favorability is modulated by subtle changes to the inter-filament rotation angle. To test this hypothesis, we examined all minima within angular shift ranges that satisfy *τ* to form fascin bonds (yellow in Figure S4L). Consistently, when the rotation angle of filament *A* is > than that of filament *B* (upper diagonal in Figure S4I), the *E_c_* minima correspond to “down” pose bridges (Figure S4M), while when the rotation angle of filament *B* is > than that of filament *A* (lower diagonal in Figure S4I), the *E_c_* minima correspond to “up” pose bridges (Figure S4N), with adjacent minima corresponding to bridging of adjacent axially-shifted protomers.

## Graphics and additional analysis

Figures and movies were generated with UCSF Chimera and ChimeraX. All statistical analysis and plotting was performed in GraphPad Prism. Custom code for tomogram denoising and downstream analysis was generated with the assistance of ChatGPT 4.0.

## References

1. Revenu, C., Athman, R., Robine, S., and Louvard, D. (2004). The co-workers of actin filaments: from cell structures to signals. Nat Rev Mol Cell Biol 5, 635–646. 10.1038/nrm1437.

2. Blanchoin, L., Boujemaa-Paterski, R., Sykes, C., and Plastino, J. (2014). Actin dynamics, architecture, and mechanics in cell motility. Physiol Rev 94, 235–263. 10.1152/physrev.00018.2013.

3. Pollard, T.D., and Cooper, J.A. (2009). Actin, a central player in cell shape and movement. Science 326, 1208–1212. 10.1126/science.1175862.

4. Chhabra, E.S., and Higgs, H.N. (2007). The many faces of actin: matching assembly factors with cellular structures. Nat Cell Biol 9, 1110–1121. 10.1038/ncb1007-1110.

5. DeRosier, D.J., Tilney, L.G., and Egelman, E. (1980). Actin in the inner ear: the remarkable structure of the stereocilium. Nature 287, 291–296. 10.1038/287291a0.

6. Tilney, L.G., Derosier, D.J., and Mulroy, M.J. (1980). The organization of actin filaments in the stereocilia of cochlear hair cells. J Cell Biol 86, 244–259. 10.1083/jcb.86.1.244.

7. Rajan, S., Kudryashov, D.S., and Reisler, E. (2023). Actin Bundles Dynamics and Architecture. Biomolecules 13, 450. 10.3390/biom13030450.

8. Mattila, P.K., and Lappalainen, P. (2008). Filopodia: molecular architecture and cellular functions. Nat Rev Mol Cell Biol 9, 446–454. 10.1038/nrm2406.

9. Sauvanet, C., Wayt, J., Pelaseyed, T., and Bretscher, A. (2015). Structure, regulation, and functional diversity of microvilli on the apical domain of epithelial cells. Annu Rev Cell Dev Biol 31, 593–621. 10.1146/annurev-cellbio-100814-125234.

10. Chan, C.E., and Odde, D.J. (2008). Traction dynamics of filopodia on compliant substrates. Science 322, 1687–1691. 10.1126/science.1163595.

11. Blake, T.C.A., and Gallop, J.L. (2023). Filopodia In Vitro and In Vivo. Annu Rev Cell Dev Biol 39, 307–329. 10.1146/annurev-cellbio-020223-025210.

12. Gupton, S.L., and Gertler, F.B. (2007). Filopodia: the fingers that do the walking. Sci STKE 2007, re5. 10.1126/stke.4002007re5.

13. Wong, S., Guo, W.-H., and Wang, Y.-L. (2014). Fibroblasts probe substrate rigidity with filopodia extensions before occupying an area. Proc Natl Acad Sci U S A 111, 17176–17181. 10.1073/pnas.1412285111.

14. Davenport, R.W., Dou, P., Rehder, V., and Kater, S.B. (1993). A sensory role for neuronal growth cone filopodia. Nature 361, 721–724. 10.1038/361721a0.

15. Dent, E.W., Kwiatkowski, A.V., Mebane, L.M., Philippar, U., Barzik, M., Rubinson, D.A., Gupton, S., Van Veen, J.E., Furman, C., Zhang, J., et al. (2007). Filopodia are required for cortical neurite initiation. Nat Cell Biol 9, 1347–1359. 10.1038/ncb1654.

16. Wood, W., Jacinto, A., Grose, R., Woolner, S., Gale, J., Wilson, C., and Martin, P. (2002). Wound healing recapitulates morphogenesis in Drosophila embryos. Nat Cell Biol 4, 907–912. 10.1038/ncb875.

17. Schaefer, A.W., Kabir, N., and Forscher, P. (2002). Filopodia and actin arcs guide the assembly and transport of two populations of microtubules with unique dynamic parameters in neuronal growth cones. J Cell Biol 158, 139–152. 10.1083/jcb.200203038.

18. Sanders, T.A., Llagostera, E., and Barna, M. (2013). Specialized filopodia direct long-range transport of SHH during vertebrate tissue patterning. Nature 497, 628–632. 10.1038/nature12157.

19. Hall, E.T., Dillard, M.E., Cleverdon, E.R., Zhang, Y., Daly, C.A., Ansari, S.S., Wakefield, R., Stewart, D.P., Pruett-Miller, S.M., Lavado, A., et al. (2024). Cytoneme signaling provides essential contributions to mammalian tissue patterning. Cell 0. 10.1016/j.cell.2023.12.003.

20. Fierro-González, J.C., White, M.D., Silva, J.C., and Plachta, N. (2013). Cadherin-dependent filopodia control preimplantation embryo compaction. Nat Cell Biol 15, 1424–1433. 10.1038/ncb2875.

21. Linder, S., Cervero, P., Eddy, R., and Condeelis, J. (2023). Mechanisms and roles of podosomes and invadopodia. Nat Rev Mol Cell Biol 24, 86–106. 10.1038/s41580-022-00530-6.

22. Shibue, T., Brooks, M.W., Inan, M.F., Reinhardt, F., and Weinberg, R.A. (2012). The outgrowth of micrometastases is enabled by the formation of filopodium-like protrusions. Cancer Discov 2, 706– 721. 10.1158/2159-8290.CD-11-0239.

23. Yamaguchi, H. (2012). Pathological roles of invadopodia in cancer invasion and metastasis. Eur J Cell Biol 91, 902–907. 10.1016/j.ejcb.2012.04.005.

24. Arjonen, A., Kaukonen, R., and Ivaska, J. (2011). Filopodia and adhesion in cancer cell motility. Cell Adh Migr 5, 421–430. 10.4161/cam.5.5.17723.

25. Jacquemet, G., Hamidi, H., and Ivaska, J. (2015). Filopodia in cell adhesion, 3D migration and cancer cell invasion. Curr Opin Cell Biol 36, 23–31. 10.1016/j.ceb.2015.06.007.

26. Otto, J.J., Kane, R.E., and Bryan, J. (1979). Formation of filopodia in coelomocytes: localization of fascin, a 58,000 dalton actin cross-linking protein. Cell 17, 285–293. 10.1016/0092-8674(79)90154-5.

27. Perrin, B.J., Strandjord, D.M., Narayanan, P., Henderson, D.M., Johnson, K.R., and Ervasti, J.M. (2013). β-Actin and fascin-2 cooperate to maintain stereocilia length. J Neurosci 33, 8114–8121. 10.1523/JNEUROSCI.0238-13.2013.

28. Vignjevic, D., Kojima, S., Aratyn, Y., Danciu, O., Svitkina, T., and Borisy, G.G. (2006). Role of fascin in filopodial protrusion. J Cell Biol 174, 863–875. 10.1083/jcb.200603013.

29. Chen, L., Yang, S., Jakoncic, J., Zhang, J.J., and Huang, X.-Y. (2010). Migrastatin analogues target fascin to block tumour metastasis. Nature 464, 1062–1066. 10.1038/nature08978.

30. Sedeh, R.S., Fedorov, A.A., Fedorov, E.V., Ono, S., Matsumura, F., Almo, S.C., and Bathe, M. (2010). Structure, evolutionary conservation, and conformational dynamics of Homo sapiens fascin-1, an F-actin crosslinking protein. J Mol Biol 400, 589–604. 10.1016/j.jmb.2010.04.043.

31. Jansen, S., Collins, A., Yang, C., Rebowski, G., Svitkina, T., and Dominguez, R. (2011). Mechanism of actin filament bundling by fascin. J Biol Chem 286, 30087–30096. 10.1074/jbc.M111.251439.

32. Yang, S., Huang, F.-K., Huang, J., Chen, S., Jakoncic, J., Leo-Macias, A., Diaz-Avalos, R., Chen, L., Zhang, J.J., and Huang, X.-Y. (2013). Molecular mechanism of fascin function in filopodial formation. J Biol Chem 288, 274–284. 10.1074/jbc.M112.427971.

33. Li, A., Dawson, J.C., Forero-Vargas, M., Spence, H.J., Yu, X., König, I., Anderson, K., and Machesky, L.M. (2010). The actin-bundling protein fascin stabilizes actin in invadopodia and potentiates protrusive invasion. Curr Biol 20, 339–345. 10.1016/j.cub.2009.12.035.

34. Van Audenhove, I., Denert, M., Boucherie, C., Pieters, L., Cornelissen, M., and Gettemans, J. (2016). Fascin Rigidity and L-plastin Flexibility Cooperate in Cancer Cell Invadopodia and Filopodia. J Biol Chem 291, 9148–9160. 10.1074/jbc.M115.706937.

35. Tan, V.Y., Lewis, S.J., Adams, J.C., and Martin, R.M. (2013). Association of fascin-1 with mortality, disease progression and metastasis in carcinomas: a systematic review and meta-analysis. BMC Med 11, 52. 10.1186/1741-7015-11-52.

36. Machesky, L.M., and Li, A. (2010). Fascin: Invasive filopodia promoting metastasis. Commun Integr Biol 3, 263–270. 10.4161/cib.3.3.11556.

37. Hashimoto, Y., Kim, D.J., and Adams, J.C. (2011). The roles of fascins in health and disease. J Pathol 224, 289–300. 10.1002/path.2894.

38. Liu, H., Zhang, Y., Li, L., Cao, J., Guo, Y., Wu, Y., and Gao, W. (2021). Fascin actin-bundling protein 1 in human cancer: promising biomarker or therapeutic target? Mol Ther Oncolytics 20, 240–264. 10.1016/j.omto.2020.12.014.

39. Minn, A.J., Gupta, G.P., Siegel, P.M., Bos, P.D., Shu, W., Giri, D.D., Viale, A., Olshen, A.B., Gerald, W.L., and Massagué, J. (2005). Genes that mediate breast cancer metastasis to lung. Nature 436, 518–524. 10.1038/nature03799.

40. Bos, P.D., Zhang, X.H.-F., Nadal, C., Shu, W., Gomis, R.R., Nguyen, D.X., Minn, A.J., van de Vijver, M.J., Gerald, W.L., Foekens, J.A., et al. (2009). Genes that mediate breast cancer metastasis to the brain. Nature 459, 1005–1009. 10.1038/nature08021.

41. Yamakita, Y., Ono, S., Matsumura, F., and Yamashiro, S. (1996). Phosphorylation of human fascin inhibits its actin binding and bundling activities. J Biol Chem 271, 12632–12638. 10.1074/jbc.271.21.12632.

42. Han, S., Huang, J., Liu, B., Xing, B., Bordeleau, F., Reinhart-King, C.A., Li, W., Zhang, J.J., and Huang, X.-Y. (2016). Improving fascin inhibitors to block tumor cell migration and metastasis. Mol Oncol 10, 966–980. 10.1016/j.molonc.2016.03.006.

43. V. Chung, F. Tsai, W. Chen, D.D. Von Hoff, E.G. Garmey, J.J. Zhang, and X.Y. Huang (2022). NP-G2-044, a First-in-Class Fascin Inhibitor, Inhibits Growth and Metastasis of Gynecologic Cancers. Eur. J. Cancer 174, S34. 10.1016/S0959-8049(22)00891-7.

44. Huang, F.-K., Han, S., Xing, B., Huang, J., Liu, B., Bordeleau, F., Reinhart-King, C.A., Zhang, J.J., and Huang, X.-Y. (2015). Targeted inhibition of fascin function blocks tumour invasion and metastatic colonization. Nat Commun 6, 7465. 10.1038/ncomms8465.

45. Mogilner, A., and Rubinstein, B. (2005). The physics of filopodial protrusion. Biophys J 89, 782– 795. 10.1529/biophysj.104.056515.

46. DeRosier, D.J., and Tilney, L.G. (1982). How actin filaments pack into bundles. Cold Spring Harb Symp Quant Biol 46 *Pt* *2*, 525–540. 10.1101/sqb.1982.046.01.049.

47. Winkelman, J.D., Suarez, C., Hocky, G.M., Harker, A.J., Morganthaler, A.N., Christensen, J.R., Voth, G.A., Bartles, J.R., and Kovar, D.R. (2016). Fascin- and α-Actinin-Bundled Networks Contain Intrinsic Structural Features that Drive Protein Sorting. Curr Biol 26, 2697–2706. 10.1016/j.cub.2016.07.080.

48. Shin, H., Purdy Drew, K.R., Bartles, J.R., Wong, G.C.L., and Grason, G.M. (2009). Cooperativity and frustration in protein-mediated parallel actin bundles. Phys Rev Lett 103, 238102. 10.1103/PhysRevLett.103.238102.

49. Claessens, M.M. a. E., Semmrich, C., Ramos, L., and Bausch, A.R. (2008). Helical twist controls the thickness of F-actin bundles. Proc Natl Acad Sci U S A 105, 8819–8822. 10.1073/pnas.0711149105.

50. DeRosier, D., Mandelkow, E., and Silliman, A. (1977). Structure of actin-containing filaments from two types of non-muscle cells. J Mol Biol 113, 679–695. 10.1016/0022-2836(77)90230-3.

51. Hylton, R.K., Heebner, J.E., Grillo, M.A., and Swulius, M.T. (2022). Cofilactin filaments regulate filopodial structure and dynamics in neuronal growth cones. Nat Commun 13, 2439. 10.1038/s41467-022-30116-x.

52. DeRosier, D.J., and Censullo, R. (1981). Structure of F-actin needles from extracts of sea urchin oocytes. J Mol Biol 146, 77–99. 10.1016/0022-2836(81)90367-3.

53. Spudich, J.A., and Amos, L.A. (1979). Structure of actin filament bundles from microvilli of sea urchin eggs. J Mol Biol 129, 319–331. 10.1016/0022-2836(79)90285-7.

54. Aramaki, S., Mayanagi, K., Jin, M., Aoyama, K., and Yasunaga, T. (2016). Filopodia formation by crosslinking of F-actin with fascin in two different binding manners. Cytoskeleton (Hoboken) 73, 365–374. 10.1002/cm.21309.

55. Atherton, J., Stouffer, M., Francis, F., and Moores, C.A. (2022). Visualising the cytoskeletal machinery in neuronal growth cones using cryo-electron tomography. J Cell Sci 135, jcs259234. 10.1242/jcs.259234.

56. Tilney, L.G., Connelly, P.S., Vranich, K.A., Shaw, M.K., and Guild, G.M. (2000). Regulation of actin filament cross-linking and bundle shape in Drosophila bristles. J Cell Biol 148, 87–100. 10.1083/jcb.148.1.87.

57. Edwards, R.A., Herrera-Sosa, H., Otto, J., and Bryan, J. (1995). Cloning and expression of a murine fascin homolog from mouse brain. J Biol Chem 270, 10764–10770. 10.1074/jbc.270.18.10764.

58. Mei, L., Reynolds, M.J., Garbett, D., Gong, R., Meyer, T., and Alushin, G.M. (2022). Structural mechanism for bidirectional actin cross-linking by T-plastin. Proc Natl Acad Sci U S A 119, e2205370119. 10.1073/pnas.2205370119.

59. Huang, J., Dey, R., Wang, Y., Jakoncic, J., Kurinov, I., and Huang, X.-Y. (2018). Structural Insights into the Induced-fit Inhibition of Fascin by a Small-Molecule Inhibitor. J Mol Biol 430, 1324– 1335. 10.1016/j.jmb.2018.03.009.

60. Pednekar, D., Tendulkar, A., and Durani, S. (2009). Electrostatics-defying interaction between arginine termini as a thermodynamic driving force in protein-protein interaction. Proteins 74, 155–163. 10.1002/prot.22142.

61. Egelman, E.H., Francis, N., and DeRosier, D.J. (1982). F-actin is a helix with a random variable twist. Nature 298, 131–135. 10.1038/298131a0.

62. Galkin, V.E., Orlova, A., Schröder, G.F., and Egelman, E.H. (2010). Structural polymorphism in F-actin. Nature Structural & Molecular Biology 17, 1318–1323. 10.1038/nsmb.1930.

63. Reynolds, M.J., Hachicho, C., Carl, A.G., Gong, R., and Alushin, G.M. (2022). Bending forces and nucleotide state jointly regulate F-actin structure. Nature 611, 380–386. 10.1038/s41586-022-05366-w.

64. De La Cruz, E.M., Roland, J., McCullough, B.R., Blanchoin, L., and Martiel, J.-L. (2010). Origin of twist-bend coupling in actin filaments. Biophys J 99, 1852–1860. 10.1016/j.bpj.2010.07.009.

65. Krey, J.F., Krystofiak, E.S., Dumont, R.A., Vijayakumar, S., Choi, D., Rivero, F., Kachar, B., Jones, S.M., and Barr-Gillespie, P.G. (2016). Plastin 1 widens stereocilia by transforming actin filament packing from hexagonal to liquid. J Cell Biol 215, 467–482. 10.1083/jcb.201606036.

66. Shin, J.-B., Krey, J.F., Hassan, A., Metlagel, Z., Tauscher, A.N., Pagana, J.M., Sherman, N.E., Jeffery, E.D., Spinelli, K.J., Zhao, H., et al. (2013). Molecular architecture of the chick vestibular hair bundle. Nat Neurosci 16, 365–374. 10.1038/nn.3312.

67. Schwebach, C.L., Kudryashova, E., Agrawal, R., Zheng, W., Egelman, E.H., and Kudryashov, D.S. (2022). Allosteric regulation controls actin-bundling properties of human plastins. Nat Struct Mol Biol 29, 519–528. 10.1038/s41594-022-00771-1.

## Methods References

1. Cordingley, M.G., Register, R.B., Callahan, P.L., Garsky, V.M., and Colonno, R.J. (1989). Cleavage of small peptides in vitro by human rhinovirus 14 3C protease expressed in Escherichia coli. J Virol 63, 5037–5045. 10.1128/JVI.63.12.5037-5045.1989.

2. Gong, R., Jiang, F., Moreland, Z.G., Reynolds, M.J., de Los Reyes, S.E., Gurel, P., Shams, A., Heidings, J.B., Bowl, M.R., Bird, J.E., et al. (2022). Structural basis for tunable control of actin dynamics by myosin-15 in mechanosensory stereocilia. Sci Adv 8, eabl4733. 10.1126/sciadv.abl4733.

3. Mastronarde, D.N. (2005). Automated electron microscope tomography using robust prediction of specimen movements. J Struct Biol 152, 36–51. 10.1016/j.jsb.2005.07.007.

4. Zheng, S.Q., Palovcak, E., Armache, J.-P., Verba, K.A., Cheng, Y., and Agard, D.A. (2017). MotionCor2: anisotropic correction of beam-induced motion for improved cryo-electron microscopy. Nat Methods 14, 331–332. 10.1038/nmeth.4193.

5. Rohou, A., and Grigorieff, N. (2015). CTFFIND4: Fast and accurate defocus estimation from electron micrographs. J Struct Biol 192, 216–221. 10.1016/j.jsb.2015.08.008.

6. Mei, L., Reynolds, M.J., Garbett, D., Gong, R., Meyer, T., and Alushin, G.M. (2022). Structural mechanism for bidirectional actin cross-linking by T-plastin. Proc Natl Acad Sci U S A 119, e2205370119. 10.1073/pnas.2205370119.

7. Tang, G., Peng, L., Baldwin, P.R., Mann, D.S., Jiang, W., Rees, I., and Ludtke, S.J. (2007). EMAN2: an extensible image processing suite for electron microscopy. J Struct Biol 157, 38–46. 10.1016/j.jsb.2006.05.009.

8. Kimanius, D., Dong, L., Sharov, G., Nakane, T., and Scheres, S.H.W. (2021). New tools for automated cryo-EM single-particle analysis in RELION-4.0. Biochem J 478, 4169–4185. 10.1042/BCJ20210708.

9. Punjani, A., Rubinstein, J.L., Fleet, D.J., and Brubaker, M.A. (2017). cryoSPARC: algorithms for rapid unsupervised cryo-EM structure determination. Nat Methods 14, 290–296. 10.1038/nmeth.4169.

10. Pettersen, E.F., Goddard, T.D., Huang, C.C., Couch, G.S., Greenblatt, D.M., Meng, E.C., and Ferrin, T.E. (2004). UCSF Chimera--a visualization system for exploratory research and analysis. J Comput Chem 25, 1605–1612. 10.1002/jcc.20084.

11. Tegunov, D., and Cramer, P. (2019). Real-time cryo-electron microscopy data preprocessing with Warp. Nat Methods 16, 1146–1152. 10.1038/s41592-019-0580-y.

12. Mastronarde, D.N., and Held, S.R. (2017). Automated tilt series alignment and tomographic reconstruction in IMOD. J Struct Biol 197, 102–113. 10.1016/j.jsb.2016.07.011.

13. Ganin, Y., Ustinova, E., Ajakan, H., Germain, P., Larochelle, H., Laviolette, F., Marchand, M., and Lempitsky, V. (2015). Domain-Adversarial Training of Neural Networks. 10.48550/ARXIV.1505.07818.

14. Chen, L., Yang, S., Jakoncic, J., Zhang, J.J., and Huang, X.-Y. (2010). Migrastatin analogues target fascin to block tumour metastasis. Nature 464, 1062–1066. 10.1038/nature08978.

15. Meng, E.C., Goddard, T.D., Pettersen, E.F., Couch, G.S., Pearson, Z.J., Morris, J.H., and Ferrin, T.E. (2023). UCSF ChimeraX: Tools for structure building and analysis. Protein Sci 32, e4792. 10.1002/pro.4792.

16. Croll, T.I. (2018). ISOLDE: a physically realistic environment for model building into low-resolution electron-density maps. Acta Crystallogr D Struct Biol 74, 519–530. 10.1107/S2059798318002425.

17. Liebschner, D., Afonine, P.V., Baker, M.L., Bunkóczi, G., Chen, V.B., Croll, T.I., Hintze, B., Hung, L.W., Jain, S., McCoy, A.J., et al. (2019). Macromolecular structure determination using X-rays, neutrons and electrons: recent developments in Phenix. Acta Crystallogr D Struct Biol 75, 861–877. 10.1107/S2059798319011471.

18. Emsley, P., Lohkamp, B., Scott, W.G., and Cowtan, K. (2010). Features and development of Coot. Acta Crystallogr D Biol Crystallogr 66, 486–501. 10.1107/S0907444910007493.

19. Williams, C.J., Headd, J.J., Moriarty, N.W., Prisant, M.G., Videau, L.L., Deis, L.N., Verma, V., Keedy, D.A., Hintze, B.J., Chen, V.B., et al. (2018). MolProbity: More and better reference data for improved all-atom structure validation. Protein Sci 27, 293–315. 10.1002/pro.3330.

20. Lee, R.A., Razaz, M., and Hayward, S. (2003). The DynDom database of protein domain motions. Bioinformatics 19, 1290–1291. 10.1093/bioinformatics/btg137.

